# Multidimensional specialization and generalization are pervasive in soil prokaryotes with generalists dominating communities and specialists more central in networks

**DOI:** 10.1101/2023.03.17.533171

**Authors:** Damian J. Hernandez, Kasey N. Kiesewetter, Brianna K. Almeida, Daniel Revillini, Michelle E. Afkhami

## Abstract

Habitat specialization underpins biological processes from species distributions to speciation. However, organisms are often described as specialists or generalists based on a single niche axis, despite facing complex, multidimensional environments. Here, we analyzed 236 prokaryotic communities across the United States demonstrating for the first time that 90% of >1,200 prokaryotes followed one of two trajectories: specialization on all niche axes (multidimensional specialization) or generalization on all axes (multidimensional generalization). We then documented that this pervasive multidimensional specialization/generalization had a wide range of ecological and evolutionary consequences. First, multidimensional specialization and generalization are highly conserved with very few transitions between these two trajectories. Second, multidimensional generalists dominated communities because they were 73 times more abundant than specialists. Lastly, multidimensional specialists played important roles in community structure with ∼220% more connections in microbiome networks. These results indicate that multidimensional generalization and specialization are evolutionarily stable with multidimensional generalists supporting larger populations and multidimensional specialists playing important roles within communities likely stemming from their overrepresentation among pollutant detoxifiers and nutrient cyclers. Taken together, we demonstrate that the vast majority of soil prokaryotes are restricted to one of two multidimensional niche trajectories, multidimensional specialization or multidimensional generalization, which then has far-reaching consequences for evolutionary transitions, microbial dominance, and community roles.

**One-Sentence Summary:** Pervasive multidimensional specialization and generalization impacts evolutionary trajectories, microbial dominance, and community roles.

## Main text

In nature, organisms navigate complex environments by embracing diverse conditions (generalists) or utilizing a smaller portion of available resources/habitats (specialists). The extent to which organisms specialize or generalize is central to many ecological and evolutionary processes such as species distributions, rates of speciation, and resilience to disturbances^1, 2^. However, studies of specialization and generalization have historically focused on only a single environmental axis which overlooks the reality that organisms experience complex, heterogeneous environments that change on many axes through space and time. Omitting environmental complexity disregards relationships across niche axes and the impact they have on organisms with important implications for their ecology, evolution, and conservation. For example, the more restrictive habitat requirements of multidimensional specialists (i.e., organisms that specialize across many niche dimensions) compared to single-axis specialists could make multidimensional specialists especially susceptible to the disturbances and intensifying stress of the Anthropocene. Despite growing interest in multidimensional specialization and generalization^2–7^, there are few, if any, empirical tests of the prevalence of multidimensional specialization and generalization and their consequent effects on evolutionary trajectories, species dominance, and ecological communities.

While plant and animal studies have been important to our foundational knowledge of specialization and generalization^1, 8, 9^, investigating multidimensional niche processes in macroorganisms can be prohibitively labor intensive for even just a few species or axes^4, 5, 10^. In contrast, relatively recent advances in next-generation sequencing now allow surveys of entire, natural microbial communities (hundreds to thousands of species) across multiple environmental axes, locations, and scales with relatively low time and resource costs. Therefore, the recent rise of microbiome studies provides a promising new avenue for efficiently investigating multidimensional specialization and multidimensional generalization and their ecological and evolutionary consequences for thousands of taxa.

Here, we quantify multidimensional specialization and generalization in thousands of co-occurring microbes at local and continental scales for the first time. To do this, we analyzed the niche breadths of >1200 prokaryotes from soil microbiomes across the continental United States along environmental axes that include some of the most important abiotic factors known to shape prokaryotic soil communities (i.e. soil pH, litter depth, soil moisture, and soil temperature as well as percent soil nitrogen, percent soil carbon, and carbon/nitrogen ratio for a subset of sites)^11^. We confirm that these soil parameters are meaningful niche axes for prokaryotes in our analyses by demonstrating that they explain approximately 67% of community variation (Supplementary Methods). This large-scale investigation was made possible by the recent launch (January 2021) of the *National Ecological Observatory Network* (*NEON*), which is the National Science Foundation’s flagship ecological repository of biological, climatic, and environmental information across the continental United States and is already among the world’s largest repositories of soil microbiome data. In this study, we determined (*i*) the frequencies of multidimensional specialization (i.e., specialization across all characterized niche dimensions) and multidimensional generalization (i.e., generalization across all characterized niche dimensions) within microbial communities. We then explored the ecology and evolution of multidimensional specialists and generalists, asking (*ii*) if evolutionary transitions from multidimensional specialist to multidimensional generalist or vice-versa are more common, (*iii*) which group is dominant within microbial communities, and (*iv*) which group plays more central roles in their communities. Our study reveals a new ecological principle of prokaryotic niches in which nearly all soil taxa follow only one of two opposing trajectories – multidimensional specialization or multidimensional generalization – and highlights how constraining taxa to these two trajectories has meaningful consequences for microbial ecology and evolution.

## Bifurcating trajectories of multidimensional specialization and multidimensional generalization is a fundamental principle of soil prokaryotes

Our evaluation of 236 microbial communities from 30 sites across the United States (Supplementary Figure 1) demonstrated that multidimensional shaping of ecological niches of microbes is ubiquitous, with multidimensional generalization occurring more commonly than multidimensional specialization. We calculated niche breadths across all axes using the standard metric of “proportional similarity”, which accounts for species resource use and how common those resources are in the environment^12, 13^. Microbial taxa (“species” as identified by mapping to the GreenGenes database) displayed a bimodal distribution in niche breadth. Which we then categorize into “specialist” and “generalist” categories based on the local minima between the two peaks in niche breadth (i.e., a heuristic delineation between low and high niche breadth; see Methods for details; Figure 1e-f). Not only did ∼90% of prokaryotes (1090/1230 taxa) in the 236 communities show consistent degree of specialization or generalization across all the axes investigated, these relationships were stronger than environmental correlations among axes and robust to different analysis decisions. Specifically, we found that ∼57% of prokaryotes (697/1230 taxa) were multidimensional generalists and ∼32% were multidimensional specialists across the four main environmental axes (i.e., soil pH, moisture, temperature, and litter depth). The bimodal distribution with ∼90% of taxa being multidimensional specialist or generalists was robust to other filtering cutoffs (e.g., including taxa that occur in only 1% or 5% of samples; Supplementary File 1), and the lower number of multidimensional specialists is unlikely to result from insufficient sequencing depth because rarefaction curves in our analyses consistently plateaued (Supplementary Figure 2). Also, when using the more lenient filtering criteria of 1% and 5% occupancy, the bimodal distribution of multidimensional generalists and multidimensional specialists still holds with 73.5% and 48.1% of taxa identified as multidimensional generalists as well as 20.4% and 42.3% of taxa identified as multidimensional specialists for the 5% and 1% occupancy cut-offs, respectively. In all cases, mixed specialization/generalization was unusual with only ∼10% of taxa showing a mixture of generalization and specialization across dimensions regardless of filtering cut-off. Our results were also robust to using an alternative niche breadth metric which accounts for the similarities among habitats that taxa occupy (i.e., the range of environmental conditions; Supplementary Methods, Supplementary File 1). All of the above highlights that multidimensional generalization and multidimensional specialization are opposing niche trajectories for soil prokaryotes which supports that when taxa generalize or specialize across one niche dimension, generalization/specialization impacts most, if not all, other axes.

**Figure 1.**
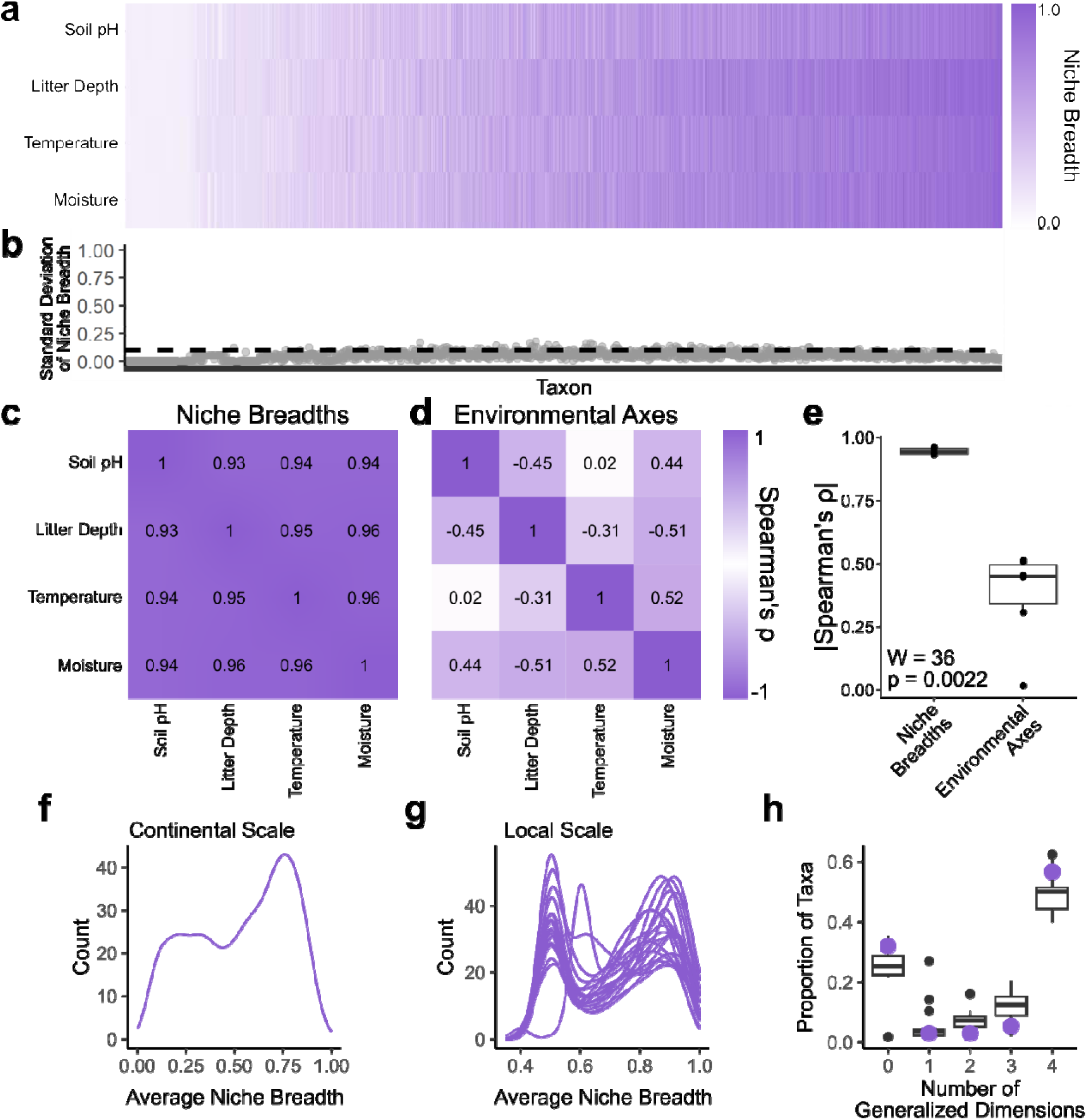
Multidimensional generalization and specialization are widespread in prokaryotes and exceed what can be explained by environmental correlations. A) Heatmap of niche breadths are highly correlated for 1230 prokaryotic taxa (x-axis) along four environmental axes (also true with seven axes; Supplementary Figure 3), indicating that most taxa (57%) are generalists across all axes and most of the remaining taxa (32%) are specialists across all axes. Taxa are ordered by average niche breadth increasing from left to right for visualization. B) Standard deviation of niche breadth across all four axes demonstrates how consistent niche breadth is across niche dimensions. Dashed line represents a standard deviation of 0.1 in proportional similarity. C) Heatmap of Spearman’s ρ from correlations between niche breadths along different axes. D) Heatmap of Spearman’s ρ from correlations between environmental axes. E) Comparison of the absolute values of Spearman’s ρ from correlations between niche breadths and correlations between environmental axes, demonstrating that niche breadth correlations are significantly stronger than correlations in environmental variation among axes. Significance determined by Mann-Whitney U test. F) Average niche breadths of taxa (mean breadth calculated across all niche dimensions) showed a bimodal distribution between specialist and generalist at a continental scale. G) Similarly, the average niche breadth of taxa calculated within each individual site also demonstrated a consistently bimodal distribution between specialist and generalist across sites. H) Distribution of multidimensional specialists/generalizations is consistent across scales. The box plot shows the distribution of taxa that generalize on n dimensions at the local scale with the purple points displaying the proportion of taxa that generalize on n dimensions at the continental scale.

In addition, microbial niche breadth on one environmental axis explained ∼80% of the variation of all other niche breadths (Spearman’s ρ = 0.94 ± 0.004; mean ± s.e.m.; Figure 1), further indicating that niche specialization and generalization occurs together along multiple environmental axes. This conclusion was supported by several additional lines of evidence. First, the relationships among niche breadths on different environmental axes when calculated at the continental scale were >10 times stronger (range: 2-54) than the correlations among environmental axes (niche breadth Spearman’s ρ = 0.90 ± 0.005 versus environmental |Spearman’s ρ| = 0.37 ± 0.078, mean ± s.e.m.; W = 36, p = 0.0022; Figure 1b-d). Second, multidimensional specialization and generalization were still ubiquitous when analyses were done at a higher taxonomic resolution with >14,000 Exact Sequence Variant taxa (Spearman’s ρ = 0.82 ± 0.010; Supplementary Figure 3a-d). Third, multidimensional specialization and generalization remained pervasive when percent carbon, percent nitrogen, and carbon:nitrogen ratios (i.e., additional major determinants of soil prokaryotic community composition available for a subset of sites) were included in our analyses (Spearman’s ρ for analysis with seven niche axes = 0.96 ± 0.001; Supplementary Figure 3e-h). Taken together, these results show that multidimensional generalization and specialization are biologically important rather than simple byproducts of correlations between environmental axes and are consistent across different taxonomic scales and different types of niche axes.

In addition to the ubiquity of multidimensional specialization and multidimensional generalization at the continental scale, it was also prevalent in soils at local geographic scales. When we determined niche breadth of taxa using the 21 individual sites with sufficient within-site replication, 75.2% ± 1.8% of taxa were specialized or generalized across all environmental axes with multidimensional generalists ∼3 times more common than multidimensional specialists, again indicating the importance of multidimensional shaping of ecological niches and that multidimensional generalization is more common than multidimensional specialization. Further, >80% of sites (17/21, t-test corrected for multiple comparisons) showed significantly stronger niche breadth relationships (5.83 ± 1.68 times greater across all sites) than environmental correlations. In short, a significant part of the multidimensional specialization and generalization is occurring independently from relationships between environmental axes (Supplementary Figure 4). Taken together, our continental and local-scale analyses emphasize that multidimensional specialization and generalization is a widespread, scale-independent ecological phenomenon in prokaryotic soil communities.

## Multidimensional generalization and specialization are bifurcating, evolutionary trajectories

Importantly, we found that this pervasive multidimensional specialization and generalization also has diverse implications for prokaryotic ecology and evolution, including for evolutionary transitions between generalist and specialist states, species dominance, and organisms’ roles within microbial networks.

First, the bifurcating trajectories of multidimensional specialists and generalists was further supported by low transition rates between specialist and generalists states, with specialist-to-generalist and generalist-to-specialist transitions occurring ∼90% less often than expected by chance. Specifically, permutational tests comparing observed transition rates to null transition distributions demonstrated that both specialist-to-generalist and generalist-specialist transitions occurred ∼90% less often than expected by chance across all 100 phylogenetic trees tested (FDR < 0.05 see Methods for details; FDR < 0.05; Figure 2a; Supplementary Figure 5). This could be due to generalization and specialization requiring specific, opposing heritable ecological strategies and adaptations. If specialization and generalization require incompatible ecological strategies/adaptations, this would allow development of generalizable frameworks for understanding prokaryotic ecology by mapping specialist/generalist strategies to other dichotomous ecological frameworks such as r/K selection theory commonly applied in microbial ecology. We find signatures of this dichotomy in the phylogenetic conservation of niche breadth with extreme specialists/generalists far more likely to have close relatives that are also as specialized/generalized (Figure 2b, see Methods for details) which is what would be expected if following specialist/generalist strategies makes it difficult to switch trajectories (i.e. are at fitness peaks on the adaptive landscape). In fact, the relationships between conservation of niche breadth and the actual value of niche breadth follows a quadratic (37.4% ± 0.02% adjusted R^2^, mean ± s.e.m.) rather than linear relationship (25.4% ± 0.01% adjusted R^2^, mean ± s.e.m.) in all 100 phylogenetic trees tested (Figure 2c; Supplementary Figure 6), indicating that the more specialized taxa are, the less likely they will evolve into generalists and vice versa. In short, multidimensional specialization and generalization are bifurcating, incompatible evolutionary trajectories.

**Figure 2.**
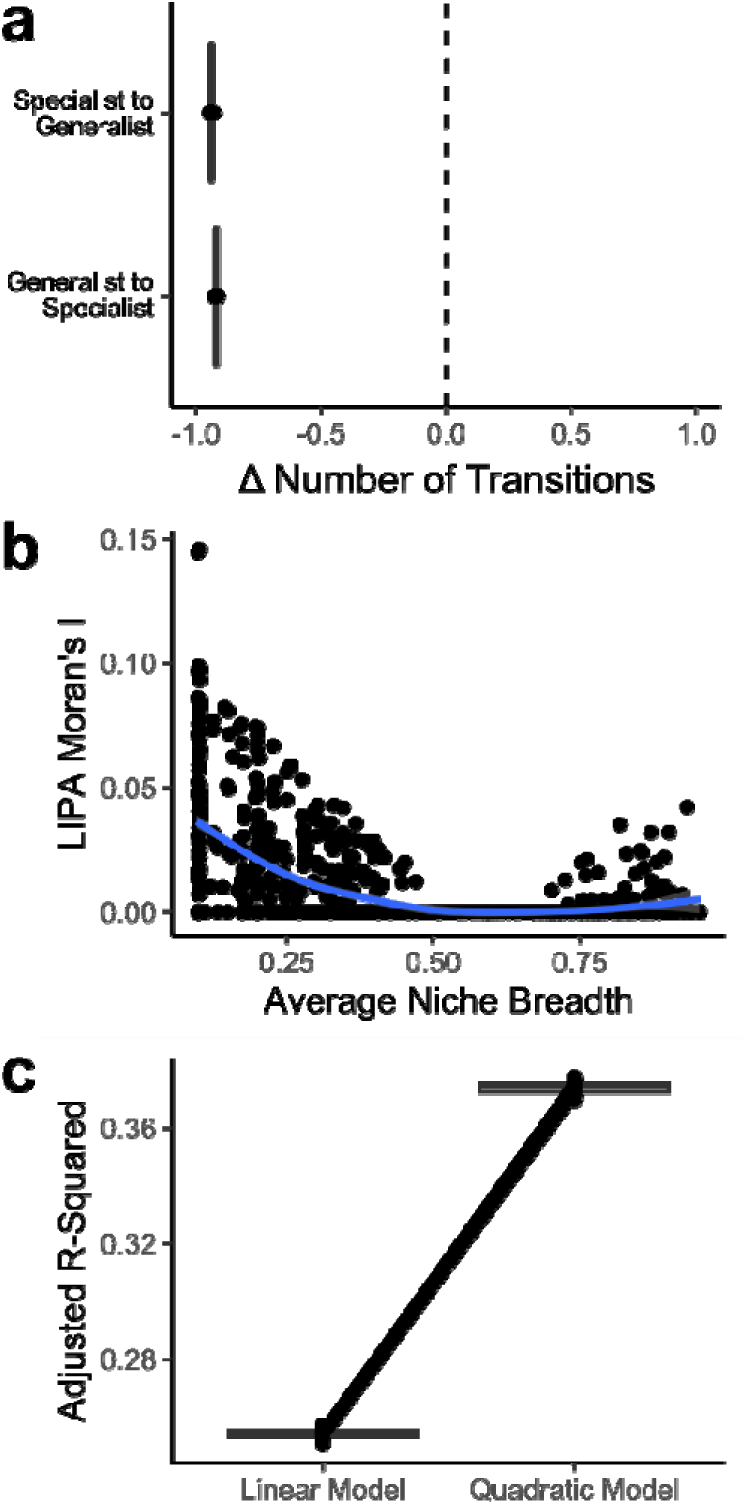
Multidimensional niche trajectories are phylogenetically conserved. A) Percent difference between number of transitions in 100 observed trees versus the null expectations (i.e., average number of transitions in the corresponding 1000 randomized trees). The dotted line indicates 0 transitions from the null expectation. Distributions of transitions in randomized trees were consistent across all 100 trees (Kolmogorov-Smirnov test) and is displayed in Supplementary Figure 5. B) LIPA Moran’s I (local measure of phylogenetic conservation of average niche breadth) in one of the 100 observed trees for all 1230 taxa (see Supplementary Figure 6 for the other 99 relationships which are all qualitatively and statistically the same as this example). The more positive the LIPA Moran’s I the more similar niche breadth is between closely related taxa. Each point represents one taxon. The blue line is a LOESS fit. Taxa with non-significant LIPA Moran’s I have an I of 0. c) Box plot comparing the adjusted R-squared values of linear and quadratic fits between average niche breadth and LIPA Moran’s I. Each line connects linear and quadratic models for the same tree. A stronger quadratic fit indicates stronger phylogenetic conservation of niche breadth at the extremes (i.e., the more generalized/specialized a taxon is, the greater the likelihood close relatives will be just as generalized/specialized).

These results suggest that multidimensional specialization and generalization may require incompatible strategies and adaptations so that becoming more specialized makes it more difficult to generalize (and vice versa). For example, shifts to a multidimensional specialist lifestyle may be difficult for generalists because specialization can require changes in multiple genes across the organisms’ genome and complex genetic regulations susceptible to mismatches in gene-gene interactions. These challenges could cause failure to specialize or to survive after specialization. Conversely, specialization may require adaptive strategies to persist that are incompatible with a generalized niche. For example, multidimensional specialists could experience a trade-off in which, instead of improving fitness by having large populations -- explored below --, they improve fitness by increasing their endurance through investing in, for example, more durable spores or other stress-tolerant traits^1, 14^ such as the endospore-forming bacteria order *Clostridiales*^15^ that is enriched for multidimensional specialists. Other obstacles could also exist that limit specialist-to-generalist transitions such as Dollo parsimony because regaining lost functions can be difficult^16–18^, multidimensional specialization could result in a loss of genes/traits essential to persisting in multiple environments thereby hindering transitions to generalist identities. This possibility is reinforced by previous work demonstrating that specialist microbes tend to have smaller genomes than generalists^19, 20^

Taken together, multidimensional generalization and specialization is an intrinsic feature of the vast majority of soil prokaryotes and likely represent two opposing evolutionary trajectories. Since multidimensional niche breadth relationships shape evolution, we then asked if multidimensional niche breadth impacts prokaryotic ecology through community dominance and structure.

## Multidimensional generalists dominate microbial communities

Second, when investigating ecological consequences of multidimensional generalization/specialization, we found that multidimensional generalists were 73 times more dominant on average than multidimensional specialists (Z = −6.28, p < 0.0001, Figure 3). Specifically, we compared the mean and maximum relative abundances (i.e., indicators of dominance/performance within communities) to determine if the size of a taxon’s niche breadth explains how dominant that taxon is relative to other taxa. Prokaryotes with wider niche breadths were more abundant than those with narrower breadths for both mean and maximum relative abundance (Spearman’s ρ = 0.84, 0.66 and p < 0.0001, p < 0.0001 for mean and maximum relative abundance, respectively; Figure 3). The dominance of generalists highlights how flexible the vast majority of the soil prokaryotic community should be to changes in environmental conditions because they can persist across a wide range of environmental conditions in many different niche dimensions.

**Figure 3.**
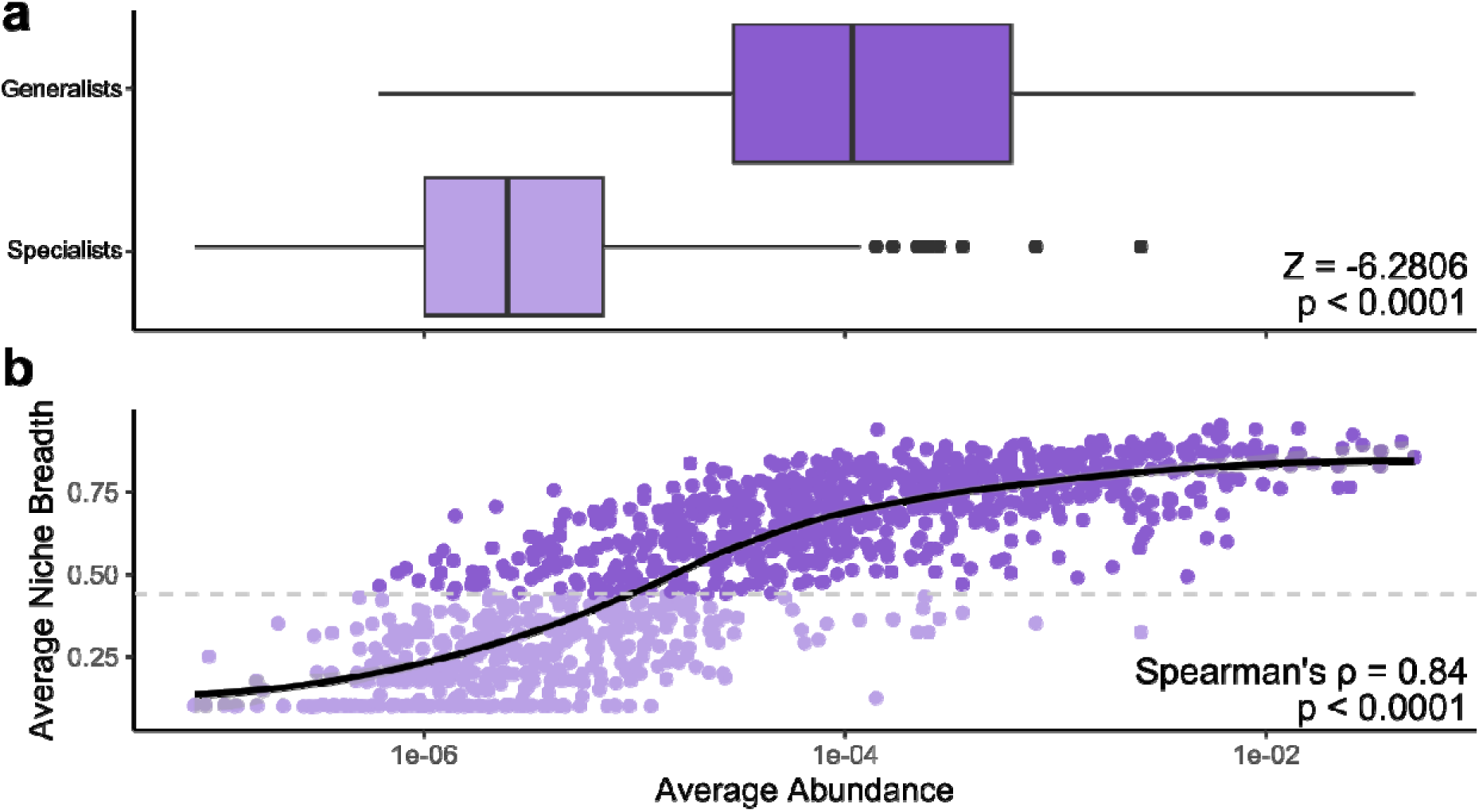
Multidimensional generalists are more dominant within communities. A) Mean abundances of generalist (dark purple) and specialist (light purple) taxa. Significance calculated with a permutational test. B) Average relative abundances of 1230 taxa regressed against average niche breadth. Line is fitted with LOESS smoothing, and shaded region around line is the 95% confidence interval. Dashed horizontal line indicates the local minima in the bimodal distribution of average niche breadth used to indicate specialists (light purple) and generalists (dark purple) in B. Direction of the relationship is determined using a Spearman’s correlation test and significance is calculated using a permutational test in which abundances are randomized 10,000 times.

The dominance of multidimensional generalists was also highly robust to a wide range of biological factors, analysis decisions, and spatial scales (Supplementary Figures 7 and 8). For instance, this relationship was maintained when analyzing communities at higher taxonomic resolutions (>14,000 Exact Sequence Variants; Spearman’s ρ = 0.62, 0.33 and p < 0.0001, p < 0.0001 for mean and maximum relative abundance, respectively; Supplementary Figure 7a,b) and when including the three additional resource axes percent carbon, percent nitrogen, and carbon:nitrogen ratios (Spearman’s ρ = 0.85, 0.70 and p < 0.0001, p < 0.0001 for mean and maximum relative abundance, respectively; Supplementary Figure 7c,d). Because dominance of multidimensional generalists could be overestimated if abundances of specialists are downweighted by absences outside of their range, we also conducted analyses accounting for sizes of each taxon’s niche breadth and the number of habitats in which a taxon is present. The goal of these two analyses was to demonstrate that our conclusions were robust when accounting for the potential of an abundance-occupancy bias. Multidimensional generalists were still dominant even after accounting for a taxon’s niche breadth (i.e., mean relative abundance/mean niche breadth) with multidimensional generalists 27 times more abundant than specialists on average (Z = 6.3857, p < 0.0001). We also found that when evaluating abundances only where taxa occur, multidimensional generalist taxa were still four times more dominant on average than specialist taxa (Z = −6.1038, p < 0.0001). Finally, the higher abundances of multidimensional generalists compared to specialists not only occurred at the continental scale, but also locally (permutational ANOVA; p < 0.0001 and p < 0.0001 for mean and maximum relative abundance; Supplementary Figure 8). In fact, niche breadth was a more important predictor of a taxon’s relative abundance than sampling site, with niche breath explaining >15 times more variation in both relative abundance metrics (⍰^2^_Niche Breadth_ = 0.52, ⍰^2^_Site_ = 0.03). Overall, our analyses indicate that multidimensional generalists are more dominant than multidimensional specialists regardless of spatial scales and taxonomic resolution, again emphasizing that most soil prokaryotes are likely to be resilient to environmental changes since multidimensional generalist persist across a wide range of conditions in many different niche dimensions and that larger population sizes may be an ecological strategy distinct to multidimensional generalists.

## Multidimensional specialists are central to community networks and functions

Finally, microbiome network analysis revealed that multidimensional specialists are more highly connected across the broader microbiome community than multidimensional generalists, indicating that multidimensional specialists are often “hub taxa” and may have keystone roles within their communities. Specifically, when we constructed microbiome co-occurrence networks for each site and assessed each taxon’s (“species”) connectedness (*i.e.,* degree centrality), specialized taxa had ∼220% greater degree centrality than generalist taxa (p < 0.0001, permutational ANOVA, Figure 4). Interestingly, specialist taxa also appear to shape the overall structure of their communities as microbiomes with greater frequencies of multidimensional specialists have significantly higher network clustering (average clustering coefficients; F_1,19_ = 13.19, p = 0.0018), making them highly-connected, tightly knit communities. Highly-connected “hub taxa” in microbiomes are often considered keystone species^21, 22^, which are organisms that play a disproportionate role in structuring communities^22–24^ which we have also recently empirically demonstrated in the field^25^. For example, the removal of a “hub” microbe in leaf endophyte and epiphyte communities destabilized communities with higher variability in community composition when the hub is absent than when it is present^21^, and highly-connected microbes (i.e. taxa with high degree centrality) shaped soil microbiome assembly in nature, repeatedly increasing biodiversity and deterministically structuring community composition during succession^25^. Therefore, while multidimensional specialists are often fewer and less abundant in communities than multidimensional generalists, their central placement within microbiome networks highlights that they likely play critical roles in these communities. Because central placement within microbiome networks and rare microbes have been documented to be important for determining microbial community variation at global scales^26^ and for supplying unique, but critical, services within communities^24, 27^, the fact that multidimensional specialists have both these characteristics and were associated with changes in microbiome-wide network properties implicates them as structurally-important taxa within communities.

**Figure 4.**
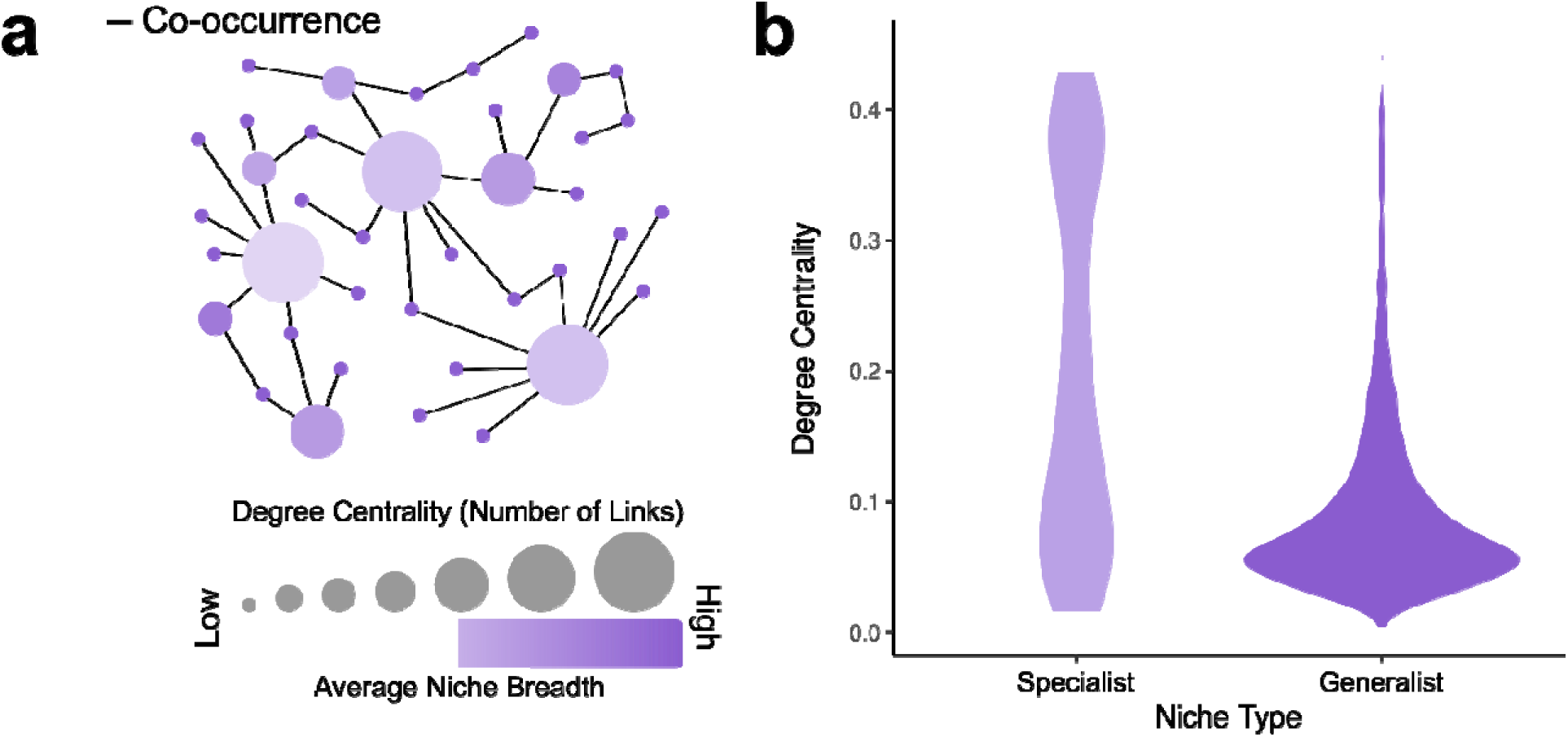
Multidimensional specialists are more central to microbiome networks. A) Schematic of analysis performed. We calculated the normalized degree centrality (number of links out of all possible links within a microbiome network) and regressed centrality against the average niche breadth of each taxon. B) Violin plot of normalized degree centrality of multidimensional specialists and multidimensional generalists. Taxa with smaller niche breadths (i.e., more specialized) are disproportionately more central than taxa with wider niche breadths (i.e., more generalized) even after accounting for site identity (p < 0.0001, permutational linear model).

Multidimensional specialists do indeed appear to play central functional roles within their communities. We found that many prokaryotic orders involved in nutrient-cycling and/or detoxification had unexpectedly high proportions of multidimensional specialists (Figure 5, Supplementary File 1) further highlighting multidimensional specialists as structurally important to their community networks. For example, 90% of *Rhodocyclales* taxa identified in our dataset are multidimensional specialists and, alongside other orders with overrepresentation of multidimensional specialists (e.g., *Desulfobacterales* and *Burkholderiales*), have been implicated in nitrogen-cycling^28^, sulfur-cycling^29^, and detoxification^29, 30^. *Clostridiales*, another order with overrepresentation of multidimensional specialists, is associated with nutrient cycling^31^ and plant symbiosis^32^. Also, many small orders of nutrient cyclers contained high numbers of multidimensional specialists; for instance, all three taxa of the green sulfur bacteria *Chlorobiales*^33^ identified in our dataset were multidimensional specialists. In essence, while multidimensional specialists may be rare, their persistence in communities and their central roles in microbial networks likely reflect their important functions in microbiomes, including crucial roles in providing nutrients and detoxifying environments. However, because the niches of these taxa are likely constrained along multiple environmental dimensions, their functions are also likely at greater threat of being lost from environmental changes than multidimensional generalists who may withstand much more variable conditions.

**Figure 5.**
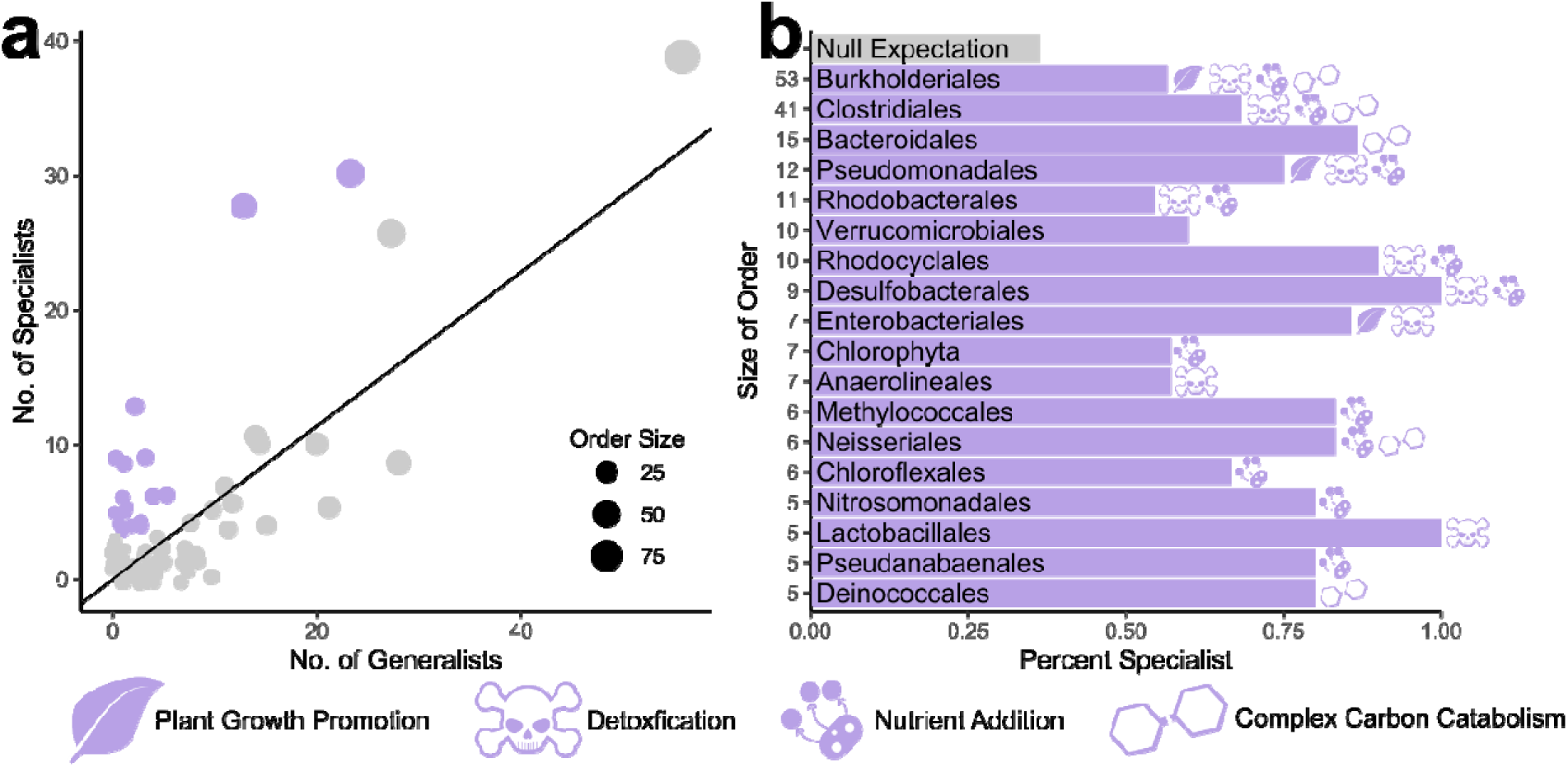
Multidimensional specialists are overrepresented in many nutrient-cycling and detoxifying orders of soil prokaryotes. A) Each dot represents an order and their sizes represent the number of taxa identified in our dataset. Purple dots are highlighted to identify orders of interest for further multidimensional specialist research (i.e., the order contained at least 5 taxa and two-fold higher ratios of multidimensional specialists to multidimensional generalists). The solid line represents the observed ratio of multidimensional specialists to multidimensional generalists across the entire dataset of prokaryotic communities, 0.571. A jitter is applied to points to better highlight the number of orders in our dataset. b) Bar chart of the purple highlighted orders in (a). The numbers on the y-axis are the sizes of the orders. The x-axis is the proportion of the orders identified as multidimensional specialist. The gray "null expectation" is the percentage of specialists in the whole dataset (36.34%). Icons to the right of columns represent functions of specialist taxa within these orders that could be verified by a literature search (literature search results provided in Supplementary File 1). "Plant growth promotion" includes bacteria that increase plant growth or improve plant defense. "Detoxification" includes heavy metal immobilization, xenobiotic degradation, hydrocarbon degradation, etc. "Nutrient Addition" includes important biogeochemical cycling processes such as carbon fixation, denitrification, sulfate reduction, nitrogen fixation, etc. "Complex Carbon Catabolism" includes breakdown of complex, difficult to metabolize carbon sources such as lignin, chitin, and cellulose. The number of multidimensional specialists and generalists for all orders are provided in Supplementary File 1.

## Conclusions

In conclusion, our study highlights multidimensional generalization and specialization as an important ecological principle in prokaryotic communities by demonstrating for the first time that virtually all soil prokaryotes follow two opposing trajectories in multidimensional niche space with cascading consequences for evolutionary transitions, taxon dominance, and microbial roles within communities. Given that these microbes undergird many ecosystem functions and services (e.g., nutrient cycling, carbon sequestration, supporting primary producers), widespread multidimensional specialization and generalization influences natural processes by shaping prokaryotic ecology and evolution. For example, ecosystem services reliant on multidimensional generalists may be more robust to environmental instability^3, 34–36^ than those reliant on multidimensional specialists since multidimensional specialists depend on maintaining a complex set of environmental conditions. The discovery of multidimensional generalization and specialization as a ubiquitous feature of prokaryote communities has also sparked many questions for future work. First, investigation into the underlying mechanisms and processes that constrain prokaryotes to those two trajectories would be profitable. In particular, we advocate for studies asking: “Do multidimensional generalists have more genetic diversity in their populations allowing them to occupy greater multidimensional niche space?”, “Can specialization on one axis restrict the environmental conditions a taxon is exposed to thereby leading to specialization across additional axes through adaptation to those restricted conditions?”, and/or “To what extent do pleiotropic and epistatic interactions among genes underpin multidimensional changes in the niche?”. While agglomerative strategies such as bulk physiological measurements (e.g., carbon flux in soil cores) or meta-genomes/meta-transcriptomes of whole soils do not allow fine enough resolution to address these types of questions, the advent of new microfluidic sequencing and culturing technologies^37–39^ could provide the fine-tuned resolution necessary to study the genetics and physiology of not just populations but the actual individual microbes that make up those populations. Additionally, some abiotic dimensions, such as oxygen availability, can vary at microhabitat levels and change dramatically along a single granule of soil^40, 41^. As a result, future work analyzing these fine-scale niche dimensions would be interesting to determine if the multidimensional relationships we found for “macrohabitat” dimensions are also important at the microhabitat scale. Microhabitat studies could also provide valuable insight into whether multidimensional specialists utilize microhabitats to avoid competitive exclusion by the community-dominating multidimensional generalists. Second, although multidimensional generalists were more dominant and somewhat more common in our study, multidimensional specialists did make up a meaningful part of the overall communities (32%) and were more often hub taxa, making their ecology especially interesting for future investigation. To understand how these multidimensional specialists may be structuring their communities, we propose that single cell sequencing approaches could be used to profile functional expression of these taxa *in situ* and microfluidic culturing approaches can be used to isolate these putative keystones for phenotyping and experimental manipulations. Third, we have analyzed multidimensional niche breadth relationships along abiotic dimensions (e.g., pH, temperature, etc.), but, moving forward, it would be valuable to also analyze biotic dimensions (e.g., host plant breadth) in order to determine if multidimensional specialization and generalization is also common for the biotic niche of prokaryotes. Unlike abiotic niche dimensions, biotic niches can be actively shaped by adaptation in the partner organisms, which may require different ecological strategies outside of multidimensional specialization and generalization. Recent work in fungi and oomycetes suggests that specialization and generalization may not be strongly correlated between abiotic and biotic niche dimensions^42–44^, but this has not been tested in soil prokaryotes. Finally, because multidimensional specialization creates more constraints on where organisms can persist, multidimensional specialists may be at greater risk from accelerating habitat loss and environmental change in the Anthropocene^10, 45^. Further, because of multidimensional specialists’ central role in their communities, the loss or decline of these taxa could perturb the entire prokaryotic community, especially in cases where they provide unique, but critical, functions. Thus, future studies testing predictions of multidimensional specialist/generalist resilience and consequent effects on ecosystem function and stability could be especially important for understanding microbial roles in ecosystem responses to global change.

## Acknowledgements

We thank the *National Ecological Observatory Network* for making their data publicly available. We also thank Amy Zanne (University of Miami), Sharon Strauss (University of California - Davis), Kerri Crawford (University of Houston), Maanasa Jayachandran (Florida International University) and Amanda Rawstern, Alexandria Igwe, and Gina Ortiz of the Afkhami lab (University of Miami) as well as the labs of Alex Wilson, Cynthia Silveira and Lena Müller (University of Miami) for their feedback on this manuscript. We also thank the three anonymous reviewers and the editor for their feedback on this manuscript.

## Funding

We acknowledge funding support from the University of Miami to DJH (Maytag Fellowship, Dean’s Summer Research Fellowship, Dean’s Dissertation Fellowship) and KNK (Lisa D. Anness Fellowship) as well as funding from the USDA to DJH (NIFA Predoctoral Fellowship 2022-67011-36456) and the National Science Foundation to KNK (Graduate Research Fellowship), BKA (Graduate Research Fellowship) and MEA (DEB-1922521 and NSF DEB-2030060).

## Author contributions

Conceptualization: DJH, KNK, MEA. Data Analyses: DJH, KNK. Data Collection: DJH, KNK, BKA, DR. Writing: DJH, KNK, MEA. Review and Editing: DJH, KNK, BKA, DR, MEA. Revisions: DJH, MEA. Supervision: MEA.

## Competing interests

All authors declare no competing interests.

## Data and material availability

All raw sequencing and environmental data are publicly available through the *NEON* database. Scripts to download data from NEON and process sequencing data into ESVs and OTUs are available in Supplementary File 2. OTU abundances from “kingdom” to “species” levels are available in Supplementary File 1. Code to replicate our analyses and a “project” folder containing all the intermediate files and statistical summaries from Rmarkdown scripts will be made fully available at Zenodo upon publication.

## SUPPLEMENTARY MATERIALS AND METHODS

All analyses and data preparation were conducted in *R* (version 4.0.2) using the packages described in Supplementary File 1 unless indicated otherwise.

### Microbial sequence and environmental data collection

For this study, we analyzed prokaryotic soil communities and environmental/biogeochemical data from 236 plots in the *National Ecological Observatory Network* (*NEON*). *NEON,* which is the National Science Foundation’s flagship ecological repository of biological, climatic, and environmental information across the continental United States, provides long-term, standardized data needed to understand ecological principles of the natural world^46^. *NEON’s* study sites are split into three hierarchical groupings (Supplementary Figure 1): site (broadest), plot, sub-plot (narrowest). *NEON* is already among the world’s largest repositories of soil microbiome data, collecting prokaryotic community and biogeochemical data from soil cores collected at each subplot which are further subdivided into “organic” and “mineral” layers (if present) and analyzed separately. Sample collection and raw data processing are described in the “NEON User Guide to Microbe Marker Gene Sequences” (DP1.10108.001; DP1.20280.001; DP1.20282.001)^47^.

To obtain data on the prokaryotic community, we downloaded raw, demultiplexed prokaryotic amplicon sequencing data from the *NEON* database^48^ using scripts in Supplementary File 2. *NEON* samples were collected from field sites at peak greenness/productivity in order to standardize across habitats. Microbial genomes were extracted by *NEON* using homogenization and lysis bead beating, and DNA was extracted using the DNEasy PowerSoil kit following the Standard Operating Procedures described^49, 50^. To survey prokaryotic communities, the hypervariable V4 region of *16S rRNA* from extracted microbial genomes was amplified using standard Earth Microbiome Project primers, *515F* and *806R*^47, 51, 52^. Amplicons were sequenced on the Illumina MiSeq platform as described in the Argonne National Laboratory (2015) and Battelle Ecology, Inc (2018) SOPs^49, 50^.

To compare prokaryotic communities and abundances in different environments, we also obtained environmental data known to greatly shape microbial communities: soil pH, soil temperature, litter depth, soil moisture, percent soil nitrogen, percent soil carbon, and carbon/nitrogen ratio; Supplementary File 1)^11, 53^. All 236 plots (30 sites) had data on soil pH, soil temperature, litter depth, and soil moisture; however, a subset of the data (84 plots, 10 sites) had additional soil chemical characteristics (percent soil carbon, percent soil nitrogen, and carbon/nitrogen ratio; Supplementary Figure 1). As a result, we analyzed niche breadth twice: once across the 236 plot dataset as well as once across the subsetted 84 plot dataset that had additional soil chemical information. Analyzing niche breadth with both the full 236 plot dataset and the subsetted 84 plot dataset allowed us to first analyze more prokaryotic communities (across fewer environmental axes) and second more environmental axes (with fewer prokaryotic communities). In the first dataset, our environmental axes explain 34% and 64% of community variation without and with spatial structure (i.e., including the site ID of the plot as a factor), respectively (db-RDA). In the second dataset, 52% and 67% of community variation is explained by the environmental axes without and with spatial structure, respectively (db-RDA). Taken together, these analyses, alongside the literature^11, 54^ demonstrate that the environmental axes we selected are important components of prokaryotic niches.

### Microbial sequence processing

To convert raw prokaryotic sequencing data to relative abundances of prokaryotes (Supplementary File 2), we processed microbial sequencing data through *QIIME2* (version 2019.1) to remove sequencing adapters and chimeras, denoise single-end reads, and classify operational taxonomic units (OTUs)^55^. In short, we denoised microbial sequencing data using *Dada2* which categorized reads into Exact Sequence Variants (ESVs)^56^. We normalized the abundances of ESVs by dividing the observed number of denoised reads for a variant by the total number of denoised reads in a sample. We further grouped ESVs into “species’’ using a Naive-Bayes classifier against the 97% taxonomy reference sequence database from *GreenGenes* (version 13.5)^57^. We constructed the classifier by using the *fit-classifier-naive-bayes* function^58^ within *QIIME2*’s *feature-classifier* plug-in on the 97% taxonomy reference database. We then used the above classifier on the ESVs by using the *classify-sklearn* function (*feature-classifier* plug-in)^55^. This allowed grouping ESVs into taxa from “kingdom” to “species” levels. We refer to “species’’ (“level 7” in *QIIME2*’s terminology) as OTUs from here onward. Sample rarefaction curves plateaued indicating that further sequencing would be unlikely to identify additional taxa (Supplementary Figure 2). To compare communities across samples, we averaged the reads for each OTU across sub-plots within a plot and repeated this for any environmental data that was also collected at the sub-plot level to avoid overrepresentation of plots that were sampled more often at the sub-plot level.

### Calculating niche breadth

We first filtered the datasets, removing OTUs that were not present in at least 10% of all plots (*i.e.,* in ≤23 of 236 plots for the larger site dataset or in ≤8 of 84 plots for the smaller site dataset). This 10% cut-off filter was used to avoid spurious niche breadths resulting from poorly represented taxa with a lack of data that prevents accurate estimation of niche breadth. Likewise, we also calculated niche breadth for taxa within a site to account for the potential that differences in niche breadth seen at the continental scale are a result of large dispersal limits or other local geographic characteristics as opposed to environmental parameters. We calculated OTU niche breadth (*NB_i_*) using proportional similarity^12^:

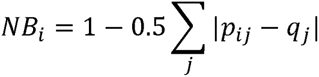

Proportional similarity constrains niche breadth between values of the *smallest q_j_* and 1 with higher values indicating wider niche breadths. Proportional similarity quantifies the habitat preference of taxa by determining if a taxon uses habitats in proportion to their availability (i.e. generalization) or disproportionately occurs within one or a few habitats (i.e., specialization). To do this, proportional similarity compares the proportion of a taxon’s population in each habitat with how common that habitat is. For instance, if the proportions of a taxon’s population mirror how common habitats are, then that taxon has no habitat preference and is generalized. Proportional similarity takes into account the proportion of taxon *i* (*p_i_*) present in habitat *j* and the proportion of all habitats that are habitat *j* (*q_j_*). Proportional similarity offers two advantages: 1) because *NB_i_* can only reach a maximum of 1, it is possible to compare niche breadths across multiple environments (this is particularly important in our analyses when niche breadth is calculated at each site) since different environments will be on the same scale and 2) proportional similarity accounts for how rare/common a habitat is in the community’s environment so that, if the proportion of a taxon’s population in a given habitat is almost equal to how common that habitat is (i.e., there is no preference), the difference between *p_ij_* and *q_j_* approaches 0 for every habitat (*j*) thus resulting in niche breadths closer to 1^12^. To identify “habitats” along our environmental axes, we binned each axis into 10 bins using the functions *cut* and *cut2* in the base and *Hmisc R* packages^59, 60^ (our results were consistent up to our highest tested number of bins, 30). We also repeated our analyses using another common niche breadth metric – niche range – which is the range between the minimum and maximum bin on any environmental axis that an organism occupies thus accounting for the adjacency of environmental bins. All our results were consistent and statistics are provided in Supplementary File 1.

### Relationships between niche breadth across axes

To determine if niche breadth is related across multiple axes (i.e., whether multidimensional niche specialization and/or generalization are common or if niche breadths have no relationship among niche axes), we correlated niche breadths of each taxon on one environmental axis (e.g., soil pH) with niche breadths on all other environmental axes using Spearman correlations. Importantly, we also examined whether multidimensional specialization and generalization were due to correlation in the environmental axes themselves (e.g., if organisms’ multidimensional generalization on soil temperature and soil moisture niche axes is because soil temperature and moisture are strongly correlated). To do this, we determined the relationship between environmental axes using Spearman correlations and compared the absolute values of those Spearman’s coefficients with the Spearman’s coefficients for the niche breadth relationships using a Mann-Whitney U test. To test whether effects were the same at local scales, we repeated these analyses at each site using niche breadths calculated at the site level with the *aovp* function (*lmPerm* package). *aovp* uses permutation tests to calculate significance in an ANOVA model (without requiring a normal distribution). This test allowed us to determine if relationships between niche breadths were stronger or weaker than relationships between environmental axes after accounting for variation associated with a sample’s originating site. To evaluate the robustness of our results to taxonomic resolution, we repeated these analyses at the higher taxonomic resolution of ESVs.

### Comparing taxon dominance of multidimensional specialists and generalists

To assess if multidimensional specialists or generalists are more dominant in communities, we determined the average and maximum relative abundances of all taxa across the 236 communities and compared differences between multidimensional specialists (all niche breadths below the local minimum in average niche breadth distributions)^13, 61^ and multidimensional generalists (all niche breadths above the local minimum) for each abundance metric with permutational tests (*independence_test* function, *coin* package). After the permutational test determined whether these specialists and generalists had different abundances, we ran Spearman’s correlations tests of the two abundance metrics against niche breadth to determine: 1) whether relative abundance increases or decreases with niche breadth and 2) how much of the variation in abundance differences could be attributed to niche breadth. We also repeated this analysis at the higher taxonomic resolution of ESVs to ensure results were robust. We conducted several other analyses to further check the results were robust. For instance, because effects on abundance may result from dispersal limits at a continental scale, we ran a similar test at the local “site” level. Specifically, we tested whether niche breadth (calculated for each taxon at each site) explained the two abundance metrics after accounting for variation associated with which site a sample originates by using the *aovp* function (*lmPerm* package). In addition, because dominance of generalists could be overestimated if abundances of specialists are down-weighted by absences outside of their range, we also conducted two analyses accounting for the potential of an abundance-occupancy bias by accounting for sizes of taxa’s niche breadths and the number of habitats in which a taxon is present.

### Calculating transition rates between multidimensional generalist and specialist taxa

To determine if multidimensional generalists and specialists are more or less likely to transition to the opposite state (e.g., generalist transitions to specialist) than expected by chance, we constructed phylogenetic trees using the microbial sequences from the *NEON* sequencing data and calculated transitions with stochastic character mapping. Because OTUs consist of multiple ESVs and observations may change based on which variant is used to represent an OTU, we constructed 100 phylogenetic trees in which a randomly chosen variant within an OTU is used to represent that OTU. We performed multiple sequence alignments using the *CLUSTAL*Ω^62^ algorithm with default parameters. We then converted the alignments to distance matrices with the *dist.ml* function (*phangorn* package) and used the Jukes and Cantor 1969 (JC69) substitution model which assumes equal frequencies of nucleotides and equal mutation rates between nucleotides.^63^ We built trees using the neighbor-joining method (*NJ* function in *phangorn* package) and rooted these trees at the midpoint (*midpoint* function in *phangorn* package)^64–66^. For each of the 100 constructed trees, we compared the observed transition rates from specialist to generalist (and vice versa) under an “Equal Rates” model using the *make.simmap* function (*phytools* package) against the transition rates of trees in which “specialist” and “generalist” status were randomized 1000 times without replacement. This allowed us to determine: 1) whether the observed transition rates in the 100 observed trees were different than expected by random chance and 2) how much the observed transition rates changed compared to random expectations. We used an “Equal Rates” model, a model which assumes that transitions between specialist and generalist status (and vice versa) occur at equal rates^67^. We calculated one-tailed p-values for the observed number of transitions from generalist to specialist status and vice versa for each of the 100 observed trees by calculating the number of transitions in 1000 randomized trees that were less than the number of transitions in the observed tree and then dividing that sum by 1000 (i.e., the number of permutations). We then corrected for multiple comparisons by calculating the False Discovery Rate (Benjamini-Hochberg correction).

To measure whether more specialized/generalized taxa have more specialized/generalized relatives than intermediately specialized/generalized taxa, we calculated Local Indicators of Phylogenetic Association (LIPA) Moran’s I of average niche breadth across all 100 previously constructed trees. LIPA Moran’s I is the same formula for Local Indicators of Spatial Association, but, instead of being applied on spatial distances, it is applied to phylogenetic distances^68^. For each taxon, a LIPA Moran’s I can be calculated for its average niche breadth to quantify if niche breadth in that area of the tree is a hotspot of phylogenetic clustering in niche breadth by applying a weighting constant of d *^-^*^1^ where *d* is the phylogenetic distance between focal taxon (*i*) to all other taxa in the phylogeny (*j*). We then determined if Local Moran’s I (i.e., niche breadth conservation) is constant throughout the entire phylogeny or highest at the two niche breadth extremes by regressing Local Moran’s I against average niche breadth. We then compared two generalized linear models, one in which Local Moran’s I was the response variable and average niche breadth was the predictor variable and a quadratic model with the same variables and an additional predictor variable of average niche breadth squared.

### Assessing relationships between niche breadth and network centrality

To determine if multidimensional specialists or generalists have central roles in their ecological communities, we used network theory to assess whether specialist or generalist taxa are more highly connected to other taxa within their microbiome community network and thus more likely to be hub taxa within microbiomes. We constructed co-occurrence networks using the *FastSpar* package^69^ for each site with at least 10 plots (21 *NEON* sites spread across the US) and removed taxa (“species”) that did not occur in at least 10% of sites (283 ± 13.7 taxa included per network, mean ± s.e.m.). *FastSpar* is an optimized reimplementation of the *SparCC* algorithm which infers correlations between taxa while limiting the occurrence of spurious correlations inherent in analyses of compositional datasets (e.g., relative abundances of taxa in communities)^70^. Links within co-occurrence networks are significant correlations between taxa abundances (as identified by *FastSpar*) which can represent interactions between connected taxa and/or shared habitat preferences^71–74^. As a result, analyzing the number of links a taxon has with others in a network (i.e., degree centrality) has been used in the literature to provide information about if a taxon is a keystone species and/or a habitat indicator species^21, 30, 75–77^. We calculated the degree centrality of each taxon in each network using the *degree_centrality* function in the *networkx* package (*Python*). The *degree_centrality* function normalizes the number of links a taxon has by the number of other taxa in a network 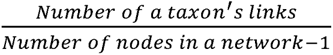 which allows for comparing degree centralities of taxa across networks of varying sizes. To identify relationships between specialization and degree centrality, we regressed the degree centrality of taxa against their average niche breadth at that site and blocked by the site from which degree centrality and average niche breadth are collected. We perform this regression using the permutational strategy in the *aovp* function (*lmPerm* package). We also analyzed global network structure of the networks and regressed average clustering coefficient (i.e., how tightly knit is the network) with the proportion of the taxa in the network that are specialists using a generalized linear model.

## SUPPLEMENTARY FIGURES

**Supplementary Figure 1.**
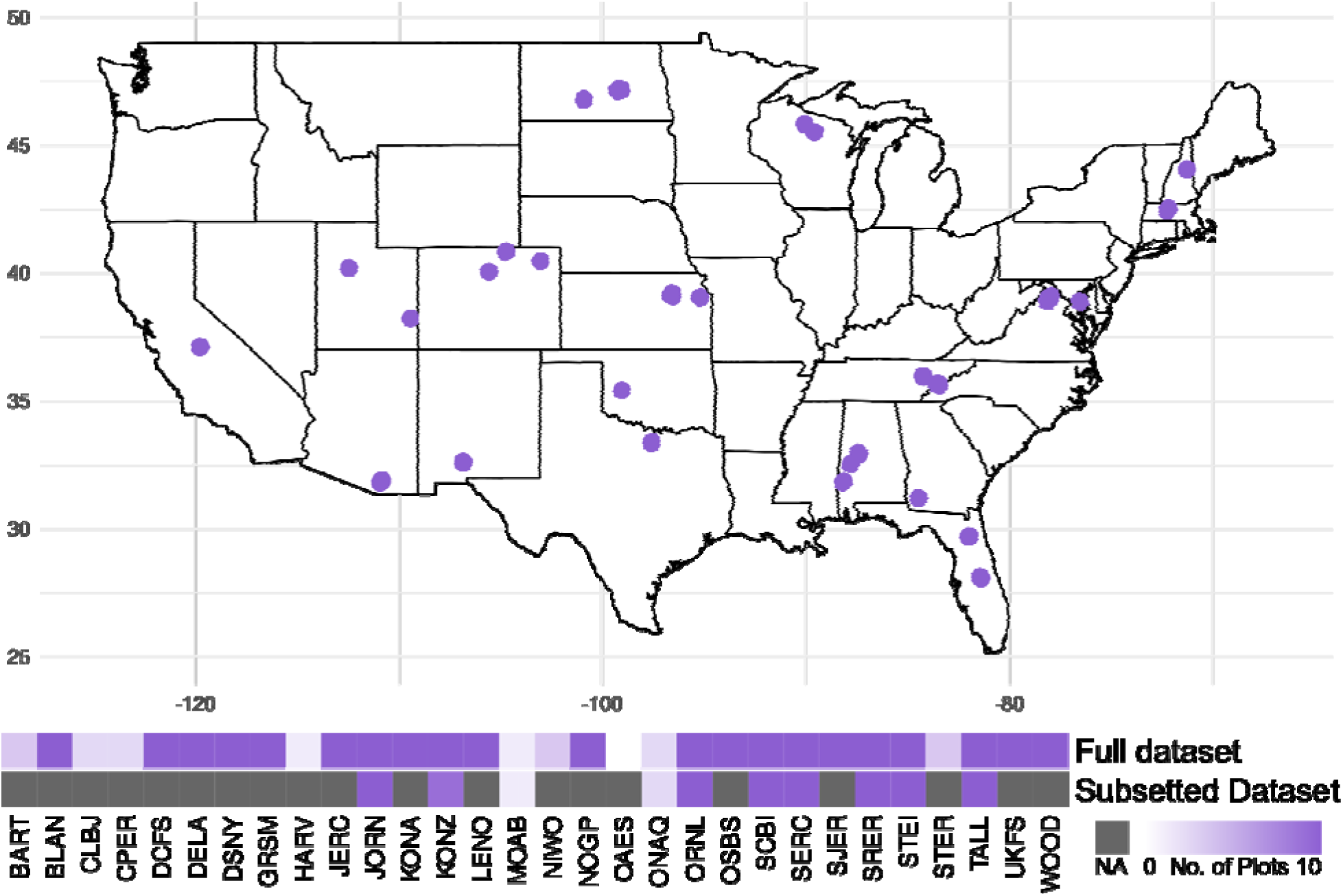
Sampling design of NEON soil collections. Map of 30 NEON collection sites across the continental United States. There are 236 plots (up to 10 plots per site) in which complete data on soil pH, soil temperature, litter depth, and soil moisture are collected (“full dataset”). A subset of the 236 plots (84 plots across 10 sites) had additional biogeochemical data on percent carbon, percent nitrogen, and carbon/nitrogen ratios (“subsetted dataset”). Distribution of the number of plots at each site is displayed in the heatmap for both the full and subsetted datasets. Dark gray squares indicate data was not available (NA)

**Supplementary Figure 2.**
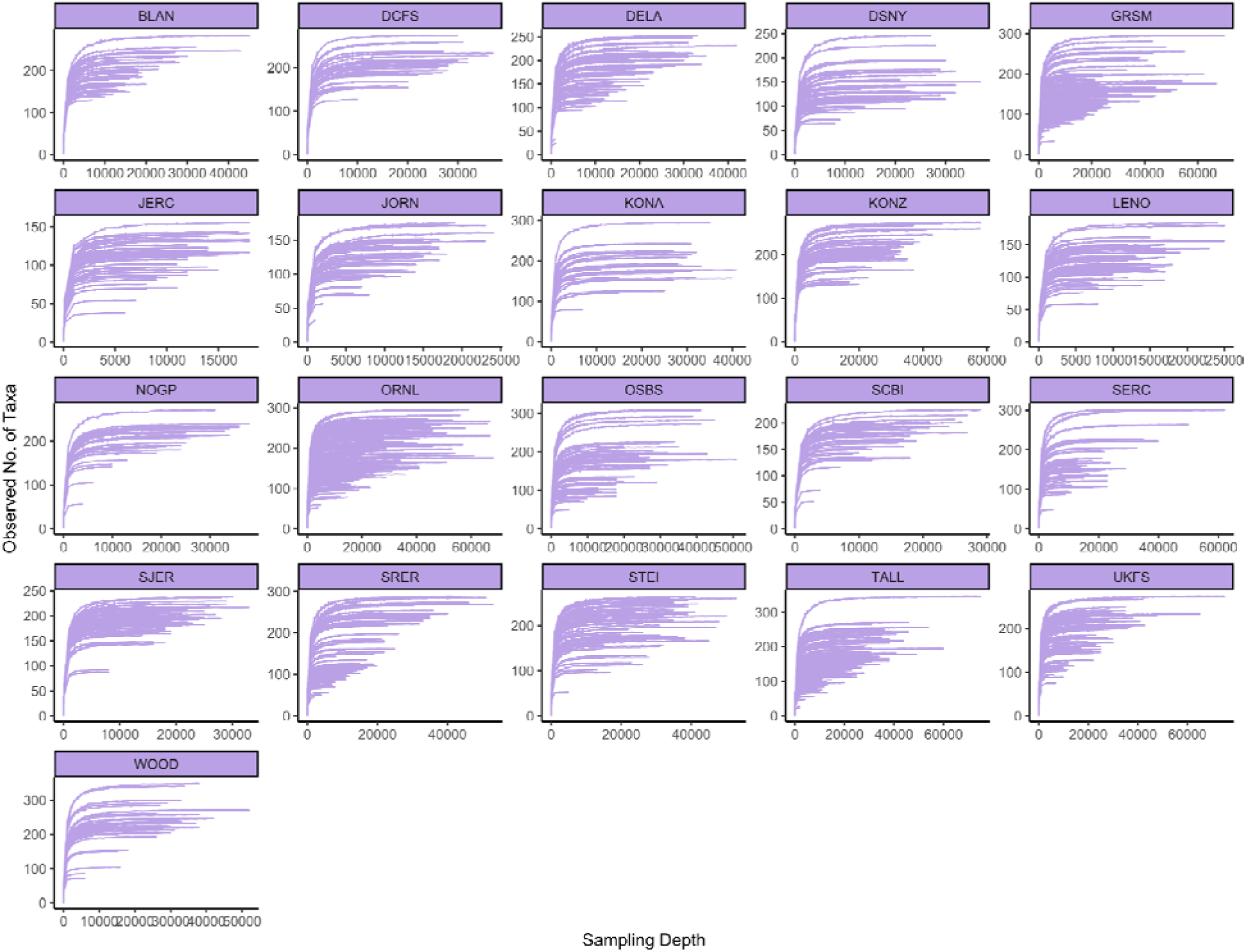
Rarefaction curves of each sample split by site. All samples reached plateaus in their rarefaction curves indicating that we had enough sequencing depth to fully characterize communities. Each line represents the rarefaction curve of a sample.

**Supplementary Figure 3.**
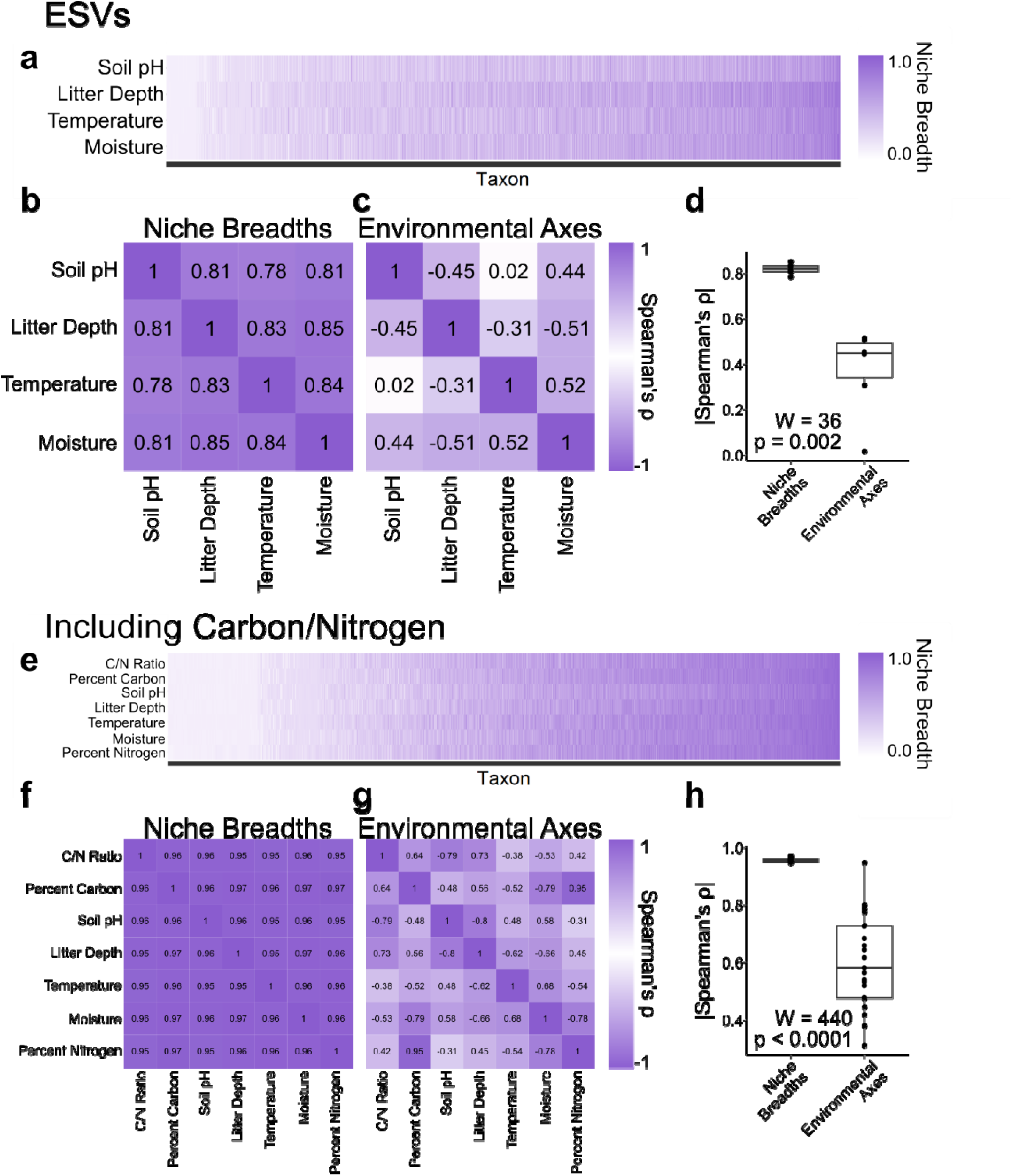
Niche breadth and environmental correlations across niche dimensions for Exact Sequence Variants (A-D) and including carbon/nitrogen niche axes (E-H) ESV analyses are represented by subfigures A-D and taxa level analyses including carbon/nitrogen data are represented by subfigures E-H. A) Heatmap of 14,015 prokaryotic ESV taxa (x-axis) along environmental axes. ESVs are sorted from lowest to highest average niche breadth for visualization. B) Heatmap of Spearman’s ρ from correlations between niche breadths of ESVs along different axes. C) Heatmap of Spearman’s ρ from correlations between environmental axes. D) Comparison of the absolute values of Spearman’s ρ from correlations between niche breadths and correlations between environmental axes, demonstrating that niche breadth correlations are significantly stronger than correlations in environmental variation among axes. Significance determined by Mann-Whitney U test. E) Heatmap of 1085 prokaryotic taxa (x-axis) along seven environmental axes that include measures of carbon and nitrogen. Taxa are sorted from lowest to highest average niche breadth for visualization. F) Heatmap of Spearman’s ρ from correlations between niche breadths along the seven different axes. G) Heatmap of Spearman’s ρ from correlations between the seven environmental axes. H) Comparison of the absolute values of Spearman’s ρ from correlations between niche breadths and correlations between environmental axes, again demonstrating that niche breadth correlations are significantly stronger than correlations in environmental variation among axes. Significance determined by Mann-Whitney U test.

**Supplementary Figure 4.**
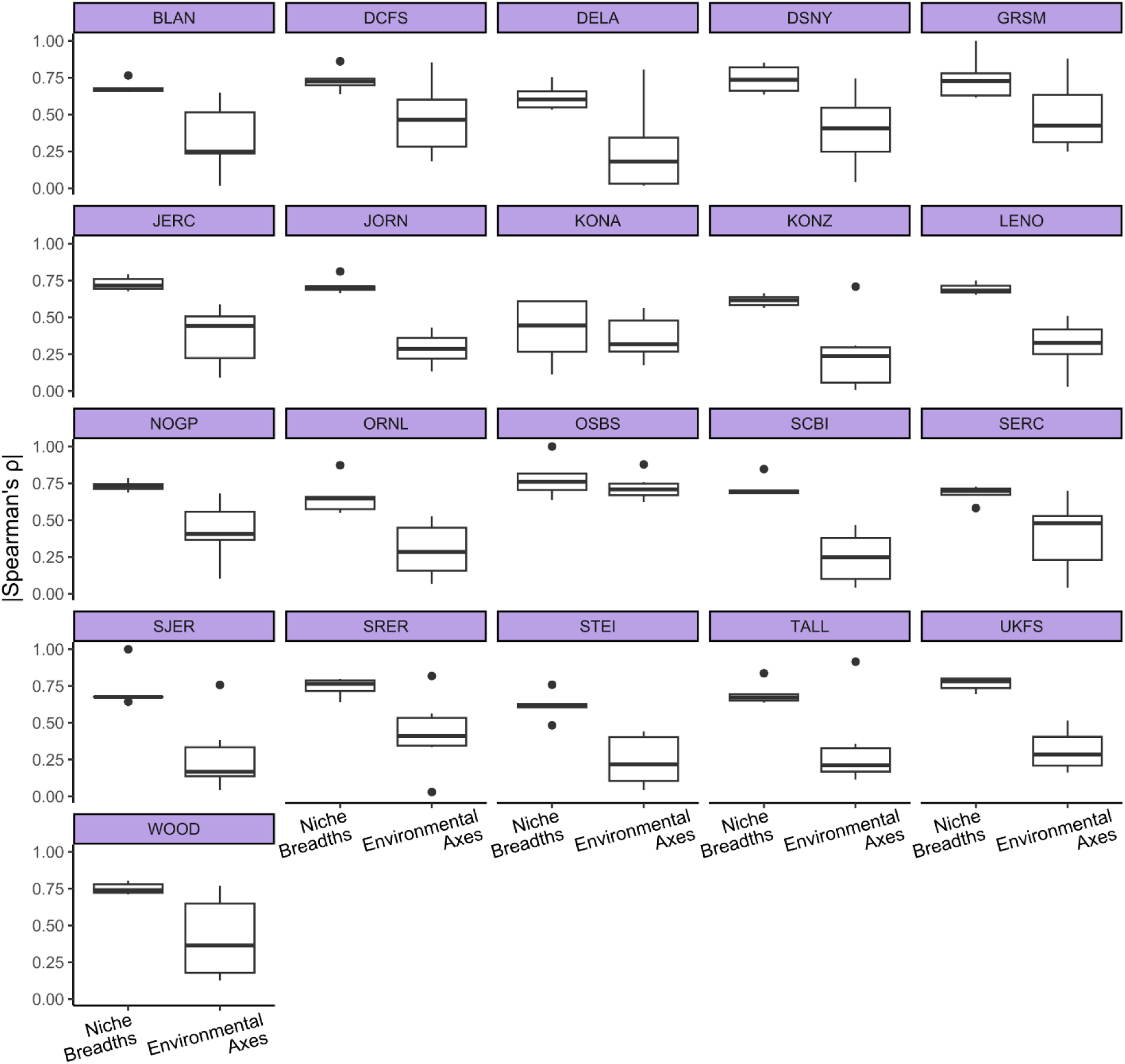
Differences in magnitude of correlations between niche breadths and environmental axes. Boxplots of the magnitude of Spearman’s coefficients between niche breadths and environmental axes at each of the 21 sites. When we account for the site from which data was collected, niche breadth relationships are still substantially stronger than environmental correlations (p < 0.0001, permutational ANOVA accounting for origin site) with the type of the relationship (i.e., relationship between niche breadths versus relationship between environmental axes) having an effect size >4 times stronger than site identity (ω_correlation type_/ω_site_ = 4.17).

**Supplementary Figure 5.**
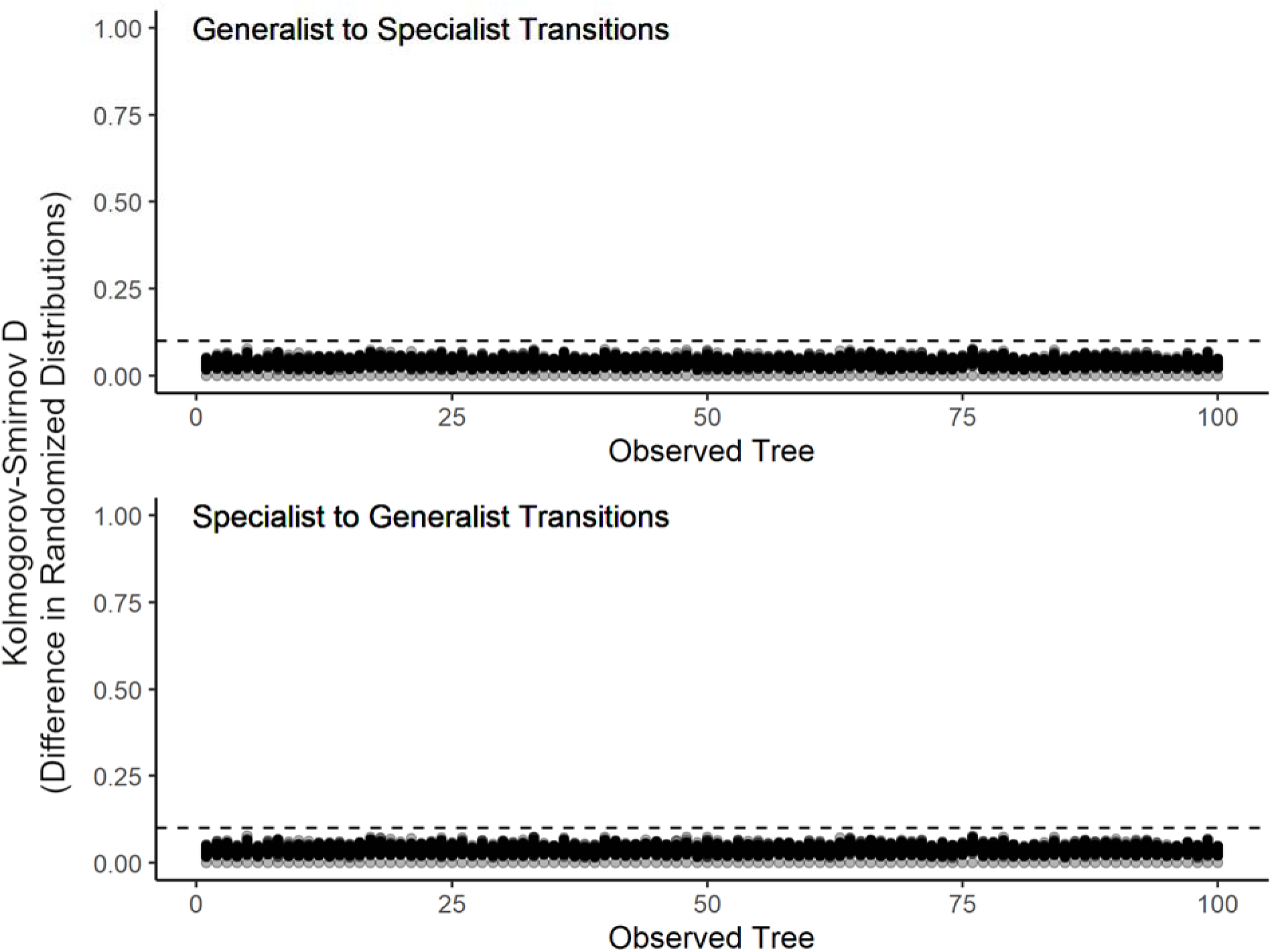
Randomized state transitions are consistent across all 100 observed representative trees. Kolomogorov-Smirnov statistics (a measure of how different the shape of two distributions are) of generalist-to-specialist (a) and specialist-to-generalist (b) transitions in all 100 observed representative trees (x-axis). Each point is the comparison of the randomized distribution of the focal representative tree (value of x-axis) against each other representative tree. All values are below a D of 0.1 (dashed horizontal line) indicating that our analyses are robust to changes in which ESV represents each OTU to which the ESV corresponds. The higher the D, the more different the distributions are from each other. The lower the D, the more similar the distributions are.

**Supplementary Figure 6.**
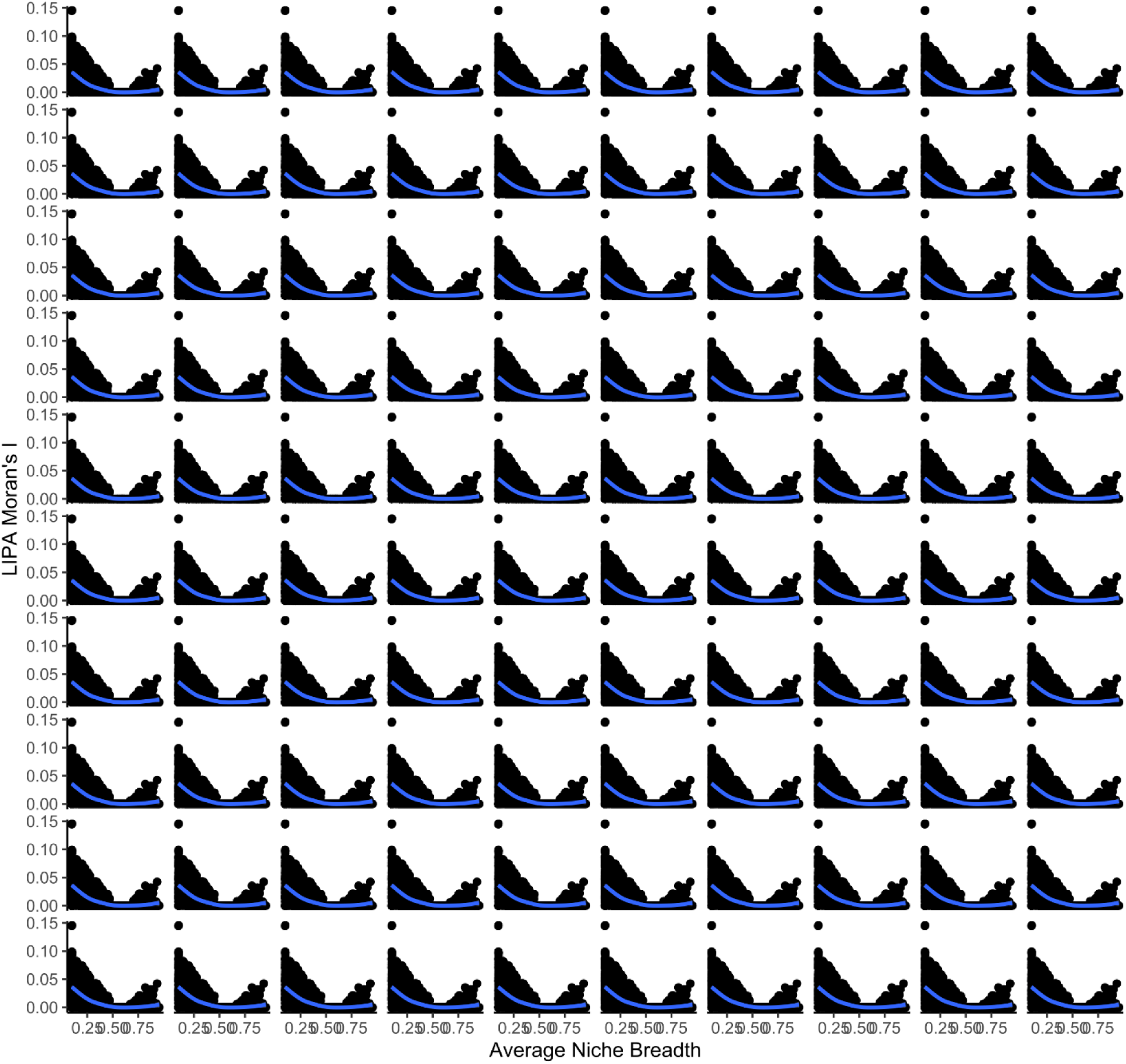
Phylogenetic relationships of niche breadth in closely-related taxa. Scatter plots of LIPA Moran’s I of average niche breadth for all 1230 taxa (points) in all 100 observed trees (each graph). A LOESS fit (blue line) is plotted to visualize if pattern follows linear or quadratic relationships (compared in Figure 2). A higher LIPA Moran’s I indicates more phylogenetic conservation of average niche breadth among closely related taxa. A quadratic relationship (a better fit than a linear model in all trees; Figure 2) indicates that phylogenetic conservation of average niche breadth is strongest when taxa are highly specialized or highly generalized. The quadratic relationship further supports multidimensional specialization and generalization as opposing niche trajectories.

**Supplementary Figure 7.**
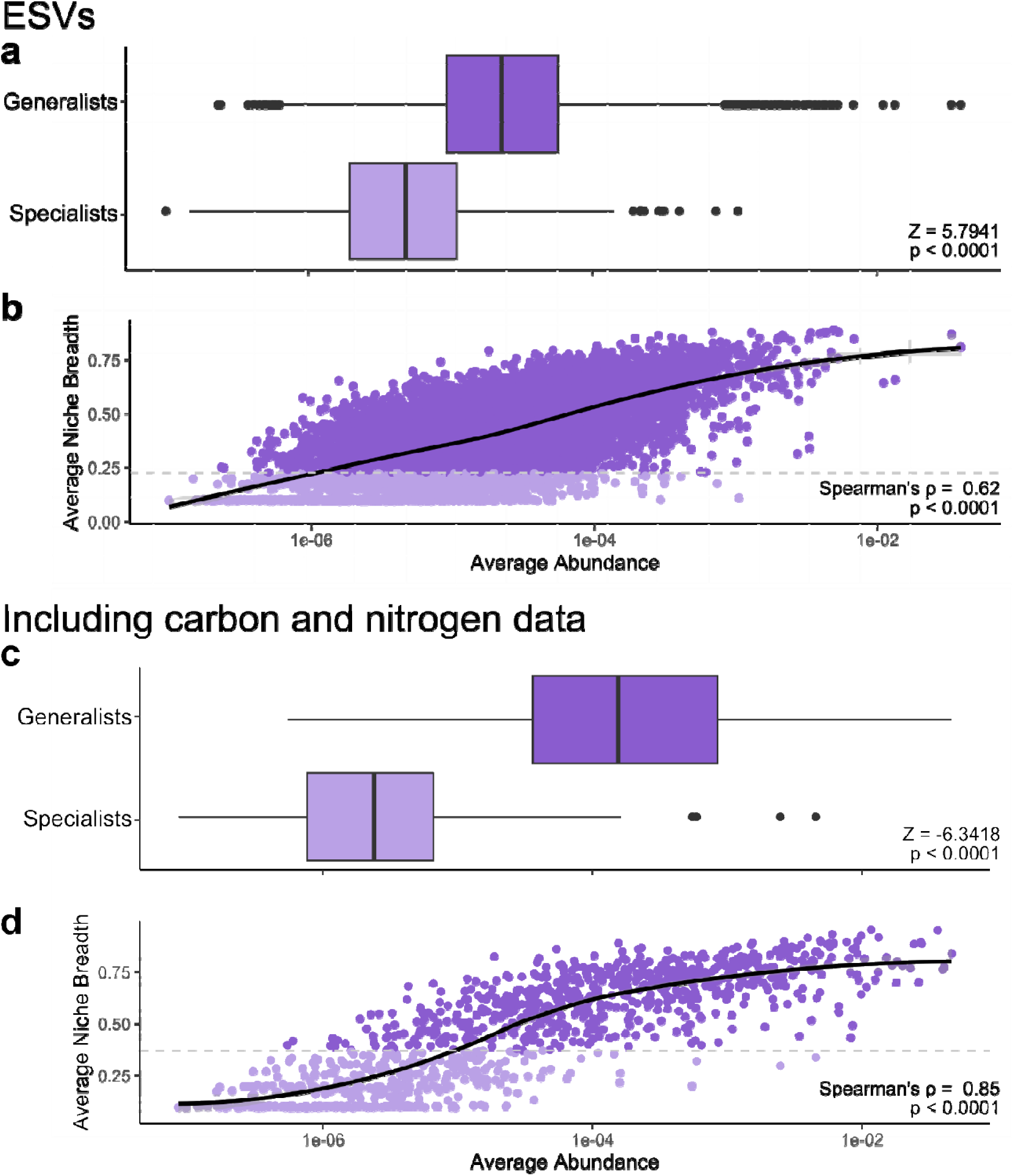
Relationship between average relative abundance and niche breadth of Exact Sequence Variants (A-B) and including carbon/nitrogen niche axes data (C-D) ESV analyses are represented by subfigures A-B and taxa level analyses including carbon/nitrogen data are represented by subfigures C-D. A) Average abundances of generalist (dark purple) and specialist (light purple) taxa. Significance calculated with a permutational test. B) Average relative abundances of 14015 prokaryotic ESV taxa regressed against average niche breadth. C) Average abundances of generalist (dark purple) and specialist (light purple) taxa. Significance calculated with a permutational test. D) Average relative abundances of 1085 taxa regressed against average niche breadth. In B and D, Lines are fitted with LOESS smoothing, shaded regions around the lines are the 95% confidence intervals, and the x-axes are on a log_10_ scale. Dashed horizontal line indicates the local minima in the bimodal distribution of average niche breadth used to indicate specialists (light purple) and generalists (dark purple). Direction of relationships were determined using a Spearman’s correlation test and significance was calculated using permutational tests in which abundances were randomized 10,000 times.

**Supplementary Figure 8.**
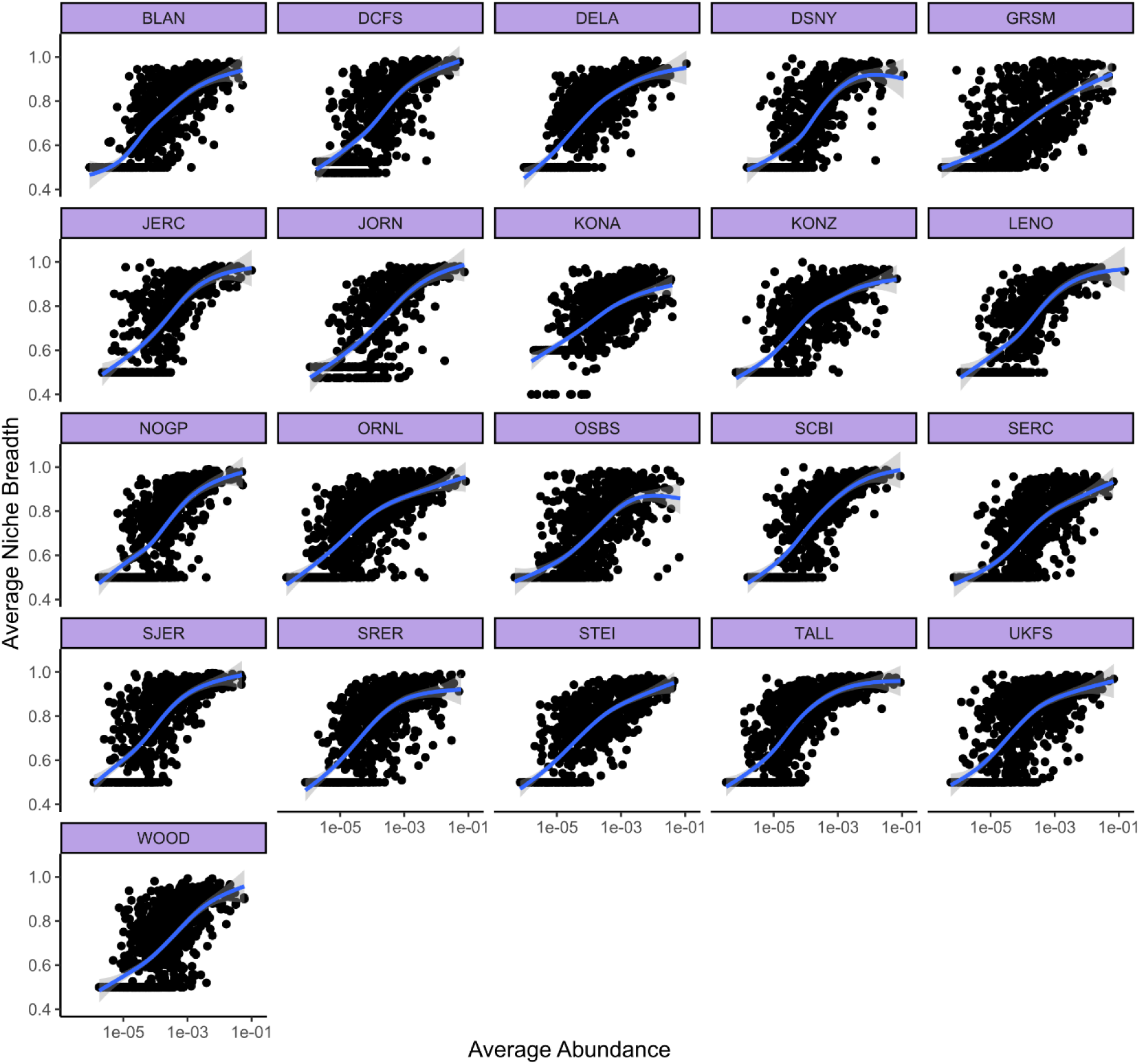
Average relative abundance within a site is explained by a taxon’s average niche breadth at that site. Average relative abundances of taxa at each of 21 sites regressed against average niche breadth in the corresponding site. Lines are fitted with LOESS smoothing, and shaded regions around the lines are the 95% confidence interval. The axes are displayed on a log_10_ scale.

## Notes

### Competing Interest Statement

The authors have declared no competing interest.

