## Supplementary File 2 for "Multidimensional specialization and generalization are pervasive in soil prokaryotes with generalists dominating communities and specialists more central in networks": Join_NEON_data_script.R

```
#For downloading NEON data and metadata

#libraries

if (!requireNamespace("BiocManager", quietly = TRUE))
  install.packages("BiocManager")
BiocManager::install()
BiocManager::install(c("GenomicRanges", "Organism.dplyr",'Epi',
                       'stringi','plyr','gtools','stringr','pheatmap',
                       'microbiome',"apeglm","forcats","DESeq2","purrr",
                       "ggthemes","lubridate","viridis","knitr",'gplots',
                       "biomformat",'phyloseq','dplyr','scales','reshape2',
                       'arm','lme4','glmm','car','plotly','outliers','GUniFrac',
                       'qiimer','devtools',"bipartite","ecodist","psych",
                       'dendextend','labdsv','effsize','miseqR','ggfortify',
                       'extrafont','picante','ape','ggplot2',
                       'ggtree','ggpubr','indicspecies','multcompView',
                       "MCMCglmm","gridExtra",'rhdf5','readr','tidyr','neonUtilities')) #rhdf5 is needed to get neonUtilities to install

install.packages("neonUtilities")
library(neonUtilities)
library(dplyr)
library(readr)
library(tidyr)

#set working directory to metadata folder
setwd("/Volumes/Macintosh HD/Users/newumuser/Desktop/")

###metadata downloads

#marker gene files for seq data 2014-2018
dir.create("neon_marker_genes_2014to2018")
zipsByProduct(dpID="DP1.10108.001", 
              startdate="2014-01", 
              enddate="2018-08",
              savepath="neon_marker_genes_2014to2018", #create folder in the correct directory on the mac, change folder for different data products w/ useful title
              package = "expanded") #this is theoretically the important part to get all the sequence data

stackByTable(filepath="neon_marker_genes_2014to2018/filesToStack10108", folder=T) #change folder title as necessary

#soil chemical properties
dir.create("soil_chem")
zipsByProduct(dpID="DP1.10078.001", 
              startdate="2014-01", 
              enddate="2018-08",
              savepath="soil_chem", #change folder for different data products w/ useful title
              package = "expanded") #this is theoretically the important part to get all the sequence data

stackByTable(filepath="soil_chem/filesToStack10078", folder=T) #change folder title as necessary

#soil physical properties
dir.create("soil_phys")
zipsByProduct(dpID="DP1.10086.001", 
              startdate="2014-01", 
              enddate="2018-08",
              savepath="soil_phys", #change folder for different data products w/ useful title
              package = "expanded") #this is theoretically the important part to get all the sequence data

stackByTable(filepath="soil_phys/filesToStack10086", folder=T) #change folder title as necessary
```
