## Supplementary File 2 for "Multidimensional specialization and generalization are pervasive in soil prokaryotes with generalists dominating communities and specialists more central in networks": qiime_16S_se.sh

```
#!/bin/sh
#This shell script contains the commands used to process the 16S V4 sequencing data obtained from NEON (Today's date 27-Aug-19).
#The assumed working directory is the root of the project folder.
#qiime2-2019.1

#Build project folder directory structure
read -p "Name your project. " project_name
read -p "Provide path to directory where you want to save this project. If you want it in this directory just type a period. " project_dir

echo Now creating ${project_name} directory in ${project_dir}.
mkdir ${project_dir}/${project_name}
cd ${project_dir}/${project_name}

#create directory tree.
echo Now creating project subdirectories in which data and analyses are stored.
mkdir -pv metadata jobs raw_seq demux_qual demux_qual/features demux_qual/visualizations phylogeny diversity diversity/visualizations taxonomy taxonomy/visualizations abundance abundance/visualizations custom_figures

read -p "Now that project directory has been constructed. Please move the sequencing data to the raw_data folder and the manifest and metadata files to the metadata folder. Once done, press any key to continue." not_real_variable

#NOTE: to view any of the visualizations (file.qzv) use the command qiime tools view path/to/file.qzv

#Input manifest data for the demultiplexed files
qiime tools import \
  --type 'SampleData[SequencesWithQuality]' \
  --input-path ./metadata/se-33-manifest_16S \
  --input-format SingleEndFastqManifestPhred33 \
  --output-path ./demux_qual/demux-single-end_16S.qza

#generate summary for demultiplexing (i.e. seqs per samples, quality distribution by position, etc.)
qiime demux summarize \
  --i-data ./demux_qual/demux-single-end_16S.qza \
  --o-visualization ./demux_qual/visualizations/demux_16S_se.qzv

qiime tools view ./demux_qual/visualizations/demux_16S_se.qzv

#Denoise reads.

qiime dada2 denoise-single \
  --i-demultiplexed-seqs ./demux_qual/demux-single-end_16S.qza \
  --p-trim-left 0 \
  --p-trunc-len 220 \
  --p-n-threads 8 \
  --o-representative-sequences ./demux_qual/rep-seqs_16S_se.qza \
  --o-table ./demux_qual/features/table_16S_se.qza \
  --o-denoising-stats ./demux_qual/stats_16S_se.qza \
  --verbose

qiime metadata tabulate \
  --m-input-file ./demux_qual/stats_16S_se.qza \
  --o-visualization ./demux_qual/visualizations/stats_16S_se.qzv

#Make directory for where to store classifiers for OTU classification.
mkdir classifiers

#Download classifiers from Greengenes (prokaryotes) and UNITE (fungi).
wget -O ./classifiers/gg_13_8_otus.tar.gz ftp://greengenes.microbio.me/greengenes_release/gg_13_5/gg_13_8_otus.tar.gz

tar -xzf ./classifiers/gg_13_8_otus.tar.gz
mv gg_13_8_otus ./classifiers

#Import classifiers for 97% homology into QIIME.
qiime tools import \
  --type FeatureData[Sequence] \
  --input-path ./classifiers/gg_13_8_otus/rep_set/97_otus.fasta \
  --output-path ./classifiers/gg_97_otus.qza

#For some reason QIIME won't read the tsv file properly for GreenGenes.
#Remove spaces from GreenGenes taxonomy file.
cat ./classifiers/gg_13_8_otus/taxonomy/97_otu_taxonomy.txt | tr -d " " > ./classifiers/gg_13_8_otus/taxonomy/97_otu_taxonomy_nospace.tsv

#Import feature reference table for taxonomic assignment.
qiime tools import \
 --type FeatureData[Taxonomy] \
 --input-path ./classifiers/gg_13_8_otus/taxonomy/97_otu_taxonomy_nospace.tsv \
 --input-format HeaderlessTSVTaxonomyFormat \
 --output-path ./classifiers/gg_97_otus_taxonomy.qza 

#Construct classifiers.
qiime feature-classifier fit-classifier-naive-bayes \
 --i-reference-reads ./classifiers/gg_97_otus.qza \
 --i-reference-taxonomy ./classifiers/gg_97_otus_taxonomy.qza \
 --o-classifier ./classifiers/gg_97-classifier-13_8.qza

#Perform classification of sequence data.
qiime feature-classifier classify-sklearn \
  --i-classifier ./classifiers/gg_97-classifier-13_8.qza \
  --i-reads ./demux_qual/rep-seqs_16S_se.qza \
  --o-classification ./taxonomy/taxonomy_16S.qza

#Visualize taxonomic assignments.
qiime metadata tabulate \
  --m-input-file ./taxonomy/taxonomy_16S.qza \
  --o-visualization ./taxonomy/visualizations/taxonomy_16S.qzv

qiime feature-table tabulate-seqs \
  --i-data ./demux_qual/rep-seqs_16S_se.qza \
  --o-visualization ./demux_qual/visualizations/rep-seqs_16S.qzv

qiime taxa barplot \
  --i-table ./demux_qual/features/table_16S_se.qza \
  --i-taxonomy ./taxonomy/taxonomy_16S.qza \
  --m-metadata-file ./metadata/sample-metadata_16S.tsv \
  --o-visualization ./taxonomy/visualizations/taxa-bar-plots_16S.qzv

qiime feature-table filter-samples \
  --i-table demux_qual/features/table_16S_se.qza \
  --p-min-frequency 2000 \
  --o-filtered-table demux_qual/features/sample-frequency-filtered-table_16S_se.qza

qiime feature-table filter-features \
  --i-table demux_qual/features/sample-frequency-filtered-table_16S_se.qza \
  --p-min-samples 10 \
  --o-filtered-table demux_qual/features/sample-contingency-filtered-table_16S_se.qza  
: <<'COMMENT'
COMMENT
```
