## Supplementary File 2 for "Multidimensional specialization and generalization are pervasive in soil prokaryotes with generalists dominating communities and specialists more central in networks": seq_download.sh

```
#Assumed working directory is root of project folder.
#The purpose of this script is to download the raw sequence files from NEON of 16S amplicon sequencing.

cd raw_data

wget https://neon-microbial-raw-seq-files.s3.data.neonscience.org/2018/BMI_BRF6G_16S_R1.fastq.tar.gz
wget https://neon-microbial-raw-seq-files.s3.data.neonscience.org/2018/BMI_BRF6G_16S_R2.fastq.tar.gz
wget https://neon-microbial-raw-seq-files.s3.data.neonscience.org/2018/BMI_C25T6_16S_R1.fastq.tar.gz
wget https://neon-microbial-raw-seq-files.s3.data.neonscience.org/2018/BMI_C25T6_16S_R2.fastq.tar.gz
wget https://neon-microbial-raw-seq-files.s3.data.neonscience.org/2018/BMI_BNMJ5_16S_R2.fastq.tar.gz
wget https://neon-microbial-raw-seq-files.s3.data.neonscience.org/2018/BMI_BNMJ5_16S_R1.fastq.tar.gz
wget https://neon-microbial-raw-seq-files.s3.data.neonscience.org/2018/BMI_C24VW_16S_R1.fastq.tar.gz
wget https://neon-microbial-raw-seq-files.s3.data.neonscience.org/2018/BMI_C24VW_16S_R2.fastq.tar.gz
wget https://neon-microbial-raw-seq-files.s3.data.neonscience.org/2018/BMI_BRPH4_16S_R2.fastq.tar.gz
wget https://neon-microbial-raw-seq-files.s3.data.neonscience.org/2018/BMI_BRPH4_16S_R1.fastq.tar.gz
wget https://neon-microbial-raw-seq-files.s3.data.neonscience.org/2018/BMI_BTV4F_16S_R2.fastq.tar.gz
wget https://neon-microbial-raw-seq-files.s3.data.neonscience.org/2018/BMI_BTV4F_16S_R1.fastq.tar.gz
wget https://neon-microbial-raw-seq-files.s3.data.neonscience.org/2018/BMI_BJ8RK_16S_R2.fastq.tar.gz
wget https://neon-microbial-raw-seq-files.s3.data.neonscience.org/2018/BMI_BJ8RK_16S_R1.fastq.tar.gz
wget https://neon-microbial-raw-seq-files.s3.data.neonscience.org/2018/BMI_BTP4N_16S_R1.fastq.tar.gz
wget https://neon-microbial-raw-seq-files.s3.data.neonscience.org/2018/BMI_BTP4N_16S_R2.fastq.tar.gz
```
